# Extrafollicular responses are sufficient to drive initiation of autoimmunity and early disease hallmarks of lupus

**DOI:** 10.1101/2022.03.04.482991

**Authors:** Lasse F. Voss, Amanda J. Howarth, Thomas R. Wittenborn, Sandra Hummelgaard, Kristian Juul-Madsen, Kristian S. Kastberg, Mathias K. Pedersen, Lisbeth Jensen, Anastasios D. Papanastasiou, Thomas Vorup-Jensen, Kathrin Weyer, Søren E. Degn

## Abstract

Many autoimmune diseases are characterized by germinal center (GC) derived affinity-matured, class-switched autoantibodies. Strategies to block GC formation and progression are currently being explored clinically, however, extrafollicular responses may also contribute to early events in autoimmunity. To investigate the relative contribution of these two pathways in autoimmune disease development, we leveraged a transgenic strategy to genetically block the GC pathway. Surprisingly, this accelerated extrafollicular responses and failed to curb autoimmune progression in two lupus models. *In vitro*, loss of the GC transcription factor Bcl-6 prevented cellular expansion and accelerated plasma cell differentiation, suggesting the *in vivo* phenotype was caused by rewiring of B cell intrinsic transcriptional programming. In a competitive scenario *in vivo*, B cells harboring the genetic GC block contributed disproportionately to the plasma cell output. Taken together, this emphasizes the extrafollicular pathway as a key contributor to autoimmune progression, and suggests that strategies aimed at blocking GCs should simultaneously target this pathway to avoid rerouting the pathogenic response.

**Highlights:**

- A genetic GC block fails to prevent autoimmune progression in two lupus models
- An intrinsic GC block drives B cell differentiation into terminally differentiated plasma cells *in vitro*
- B cells harboring a GC block competitively contribute to the plasma cell compartment in an autoreactive setting *in vivo*
- Lupus mice with a GC block display immune complex deposition in kidney glomeruli that is indistinguishable from their wild-type counterparts

**Summary:** Affinity-matured autoantibodies generated in germinal centers are a hallmark of autoimmune diseases. Voss et al. block germinal centers in two autoimmune models, but surprisingly find that disease progresses unimpeded. They identify the extrafollicular pathway as a ‘backdoor to autoimmunity’.

## Introduction

Many autoimmune diseases, such as systemic lupus erythematosus (SLE) and Sjögren’s syndrome, are characterized by the development of autoantibodies targeting nuclear antigens (Psianou et al., 2018; Rahman and Isenberg, 2008). Such antibodies can be produced by B cells via two routes, the extrafollicular pathway and the germinal center (GC) pathway (Elsner and Shlomchik, 2020). The extrafollicular pathway leads to the rapid expansion and differentiation of B cells to plasmablasts (PBs) and short-lived plasma cells (PCs), which in a natural infection setting provides rapid initial protection against pathogens. The GC pathway is slower, but enables a higher-quality response characterized by extensive somatic hypermutation and affinity maturation, robust memory cell generation, and production of long-lived PCs (Victora and Nussenzweig, 2022). At an individual B cell level, the decision between extrafollicular and GC pathway differentiation appears to rest on the initial affinity for antigen (Paus et al., 2006). While both pathways can support class-switch recombination and affinity maturation, the antibody class diversity and extent of somatic hypermutation is much greater through the GC pathway (Sweet et al., 2010; William et al., 2002).

Upon initial activation of B cells by their cognate antigen, they can either T-independently or T-dependently form an extrafollicular focus. Here they proliferate, as well as potentially class-switch, and may additionally undergo a low degree of somatic hypermutation, before they differentiate into PCs. In the context of autoimmune diseases, it has been noted that initial B cell reactivities often target autoantigenic components that carry endogenous TLR ligands capable of stimulating them independently of T cells (Lau et al., 2005; Leadbetter et al., 2002; Sweet et al., 2011). Of note, these seem strictly limited to components topologically linked to the B cell receptor (Green et al., 2021). Interestingly, signals that drive initial B cell activation also seem to limit the extent of affinity maturation (Akkaya et al., 2018). However, as autoimmune disease progresses, the breadth of the autoantigenic response often broadens, leading to inclusion of T-dependent reactivities (Cornaby et al., 2015). In both humans and mice, this broadening of the response, termed epitope spreading, has been observed even before the onset of clinical symptoms (Arbuckle et al., 2003; Degn et al., 2017b).

Epitope spreading is thought to occur through the GC pathway. In this alternate outcome of initial B cell activation, the B cells may form a primary focus in the interfollicular region, where they undergo limited proliferation and may class-switch (Roco et al., 2019; Toellner et al., 1996). They then subsequently co-migrate with cognate T cells into the follicle, where they can form a GC. In the GC, the B cells proliferate rapidly and form a ‘dark zone’, and are now termed centroblasts (Victora and Nussenzweig, 2022). The centroblasts undergo somatic hypermutation to diversify their B cell receptors. They subsequently migrate to the ‘light zone’, where they scan follicular dendritic cells for antigen. The B cells that display the highest affinity for antigen can competitively acquire antigen and present derived peptides to T follicular helper (T_FH_) cells. B cells that receive cognate T cell help may return to the dark zone for another round of division and hypermutation. B cells that do not receive help perish through programmed cell death and are engulfed by tingible body macrophages (Victora and Nussenzweig, 2022). Due to the power of the GC pathway, and the risk for inadvertent emergence of novel (auto)reactivities that are distinct from the original antigenic target, it is subject to stringent control. The requirement for T cell selection subsequent to every successive round of hypermutation, a phenomenon termed linked recognition, restricts inadvertent broadening of the response. An additional layer of control appears to be exerted by a specialized subset of T regulatory cells, termed T follicular regulatory (T_FR_) cells (Fahlquist Hagert and Degn, 2020). Nonetheless, these mechanisms appear to fail in autoimmune disease, which frequently display rampant GC activity (Domeier et al., 2017; Luzina et al., 2001).

Hence, we hypothesized that the GC pathway is critical to the autoimmune process, and that a genetic block of the GC pathway *in vivo* would prevent autoimmune development. To our surprise, a global block in the GC pathway *in vivo* in two independent models did not ameliorate autoimmune disease, but if anything exacerbated it. In an *in vitro* GC B cell culture system, GC blocked B cells expanded to a lesser extent, but were found to more rapidly develop into PBs and PCs. In a competitive scenario *in vivo*, GC blocked B cells competed efficiently with their wild-type counterparts, disproportionate to their inability to participate in GCs. Our determination of the relative contributions of the extrafollicular and GC pathways to autoimmune progression highlight a critical role of extrafollicular responses in driving autoimmune development.

## Materials and methods

### Mice

The Bcl-6^flx/flx^ strain (Hollister et al., 2013) and congenic B6.CD45.1 (B6.SJL-Ptprc^a^ Pepc^b^/BoyJ) were purchased from Jackson Laboratories (stock no. 023737 and 002014, respectively). Aicda-Cre transgenic mice (Kwon et al., 2008) were kindly provided by Meinrad Busslinger, The Research Institute of Molecular Pathology (IMP), Vienna. Aicda-Cre and Bcl-6^flx/flx^ strains were intercrossed to generate Aicda-Cre+ and Aicda-Cre− Bcl-6^flx/flx^ littermates. 564Igi mice (Berland et al., 2006) (B6.Cg-Igh^tm1(Igh564)Tik^Igk^tm1(Igk564)Tik^/J) were kindly made available by Theresa Imanishi-Kari, Tufts University, and provided by Michael C. Carroll, Boston Children’s Hospital. The 564Igi line was crossed to Aicda-Cre Bcl6^flx/flx^ to generate 564Igi H^+/+^ K^+/+^ Aicda-Cre+/- Bcl6^flx/flx^ littermates. Mice were housed in the Animal Facility at the Department of Biomedicine, Aarhus University, Denmark, under specific pathogen-free (SPF) conditions, on a 12-hour light/dark cycle with standard chow and water *ad libitum*. Both male and female mice were used in experiments. Mice were between 8 and 14 weeks old upon initiation of experiments.

### Ethics Statement

All animal experiments were conducted in accordance with the guidelines of the European Community and were approved by the Danish Animal Experiments Inspectorate (protocol numbers 2017-15-0201-01348 and 2017-15-0201-01319).

### R848 treatment protocol

Mice were treated topically on the right ear 3 times per week for 4 weeks with 1 mg R848/mL acetone using a cotton applicator or did not receive any treatment (Untreated/Unt).

### Mixed bone marrow chimeras

Recipient mice were irradiated with 9 Gy in a MultiRad 350 (Faxitron), with 350 kV, 11.4 mA, a Thoraeus filter [0.75 mm Tin (Sn), 0.25 mm Copper (Cu), and 1.5 mm Aluminium (Al)], and with a beam-distance of 37 cm. Irradiated recipients were kept on antibiotic water (either 1 mg sulfadiazine together with 0.2 mg trimethoprim per mL drinking water, or 0.25 mg amoxicillin per mL drinking water) to avoid any opportunistic infections. On the following day, donor mice were anesthetized with 4% isoflurane and euthanized. Femora, fibulae/tibiae, ossa coxae and humeri were harvested, mechanically cleaned and rinsed in FACS buffer. The Bone marrow (BM) cells were released from the harvested bones by crushing and the cell extract was then passed through a 70 µm cell strainer. The donor BM cells were then counted in a Cellometer K2 cell counter (Nexcelom). Cells were pelleted by centrifugation (200 *g*, 10 min, 4°C) and resuspended to 1*10^8^ cells/mL. Donor cells from three different mice were then mixed according to the proportions mentioned in the figure legend. The donor cell mixtures were used to reconstitute the recipient mice by retroorbital injection of 200 µL (containing a total of 20*10^6^ cells) into each recipient mouse. The reconstituted recipient mice were placed on antibiotic water the following 14 days.

### Tissue preparation

Mice were anesthetized with isoflurane (055226, ScanVet), blood samples from the retroorbital plexus were collected, and mice were euthanized using 100-150 mg/kg sodium pentobarbital (450009, Dechra Veterinary Products). Mesenteric lymph nodes (MesLN) and inguinal lymph nodes (IngLN) were removed, the splenic artery was clamped with a hemostat, and the spleen was removed. The mice were perfused intracardially with PBS (BE17-515Q, Lonza) to remove the blood, and subsequently perfused with 4% w/v paraformaldehyde (PFA) (1.04005.100, Merck) in PBS to fix the tissues. Finally, kidneys and auricular lymph nodes (AurLNs) were removed.

Collected blood samples were centrifuged at 3,000 *g* for 10 minutes, the supernatant was collected, and centrifuged again at 20,000 *g* for 3 minutes. Serum samples were stored at −20°C. The spleen and AurLNs was directly embedded in Tissue-Tek O.C.T. media (4583, Sakura Finetek) and frozen at −20°C for histology. The kidneys were kept in 4% w/v PFA for 24 hours, and then changed to 30% w/v sucrose in PBS. A small part of the spleen as well as IngLN and MesLN were stored in fluorescence-associated cell sorting (FACS) buffer (PBS, 2% heat-inactivated fetal calf serum (FCS), 1 mM ethylenediaminetetraacetic acid (EDTA)) for FACS typing.

### Flow cytometry

Spleen, IngLN and MesLN were harvested, stored into ice-cold FACS buffer, and mechanically dissociated using pestles. Spleen and LNs were filtered through 70 𝜇m cell strainers. Spleen samples were centrifuged at 200 *g* for 5 minutes at 4°C, lysed in RBC lysis buffer (155 mM NH_4_Cl, 12 mM NaHCO_3_, 0.1 mM EDTA), incubated at RT for 3 minutes, centrifuged, and finally resuspended in FACS buffer or calcium-containing buffer (PBS, 20 mM HEPES, 145 mM NaCl, 5 mM CaCl_2_, 2% FBS) when Annexin-V was included in the panels. Samples were filtered through 70 𝜇m cell strainers. Twenty 𝜇L Fc-block (553142, BD) diluted 1:50 in PBS and 100 𝜇L of each sample was added onto a 96-well plate and incubated for 5-10 minutes. Antibodies and fixable viability dye (65-0865-14, ThermoFisher Scientific) were diluted in FACS buffer or calcium-containing buffer as indicated below. One hundred 𝜇L antibody mix was added to each sample well and incubated for 30 minutes on ice. The plate was centrifuged at 200 *g* for 5 minutes, supernatant was removed, and cells were fixed for 30 minutes in PBS, 0.9% formaldehyde (F1635, Sigma-Aldrich) at RT. Later, the plates were centrifuged at 200 *g* for 5 minutes, the supernatant discarded, and the samples resuspended in FACS buffer or calcium-containing buffer. Flow cytometry evaluation was performed the following day using a 4-laser (405 nm, 488 nm, 561 nm, 640 nm) LSRFortessa analyzer (BD instruments). The following antibodies and reagents were used for flow cytometry experiments: Annexin-V-AF488 (A13201, ThermoFisher Scientific, 1:500), B220-PB clone RA3-6B2 (558108, BD, 1:500), B220-PerCP-Cy5.5 clone RA3-6B2 (561101, BD Biosciences, 1:500), CD4-PerCP clone RM4-5 (100538, BioLegend, 1:500), CD8-PerCP-Cy5.5 clone SK1 (565310, BD, 1:500), CD38-PE-Cy7 clone 90 (102718, BioLegend, 1:500), CD45.1-FITC clone A20 (110706, BioLegend, 1:500), CD45.2-APC clone 104 (109814, BioLegend, 1:500), CD95-PE clone Jo2 (554258, BD, 1:500), CD138-BV650 clone 281-2 (564068, BD, 1:500), 9D11-biotin (hybridoma kindly provided by Elisabeth Alicot, Boston Children’s Hospital, produced, purified and biotinylated in-house, 1:300), Ly6G/C-APC-R700 clone RB6-8C5 (565510, BD, 1:500), Viability Dye eFlour 780 (65-0865-14, Thermo Fisher Scientific, 1:2000), Streptavidin-BV786 (563858, BD Biosciences, 1:500), hCD2-PB clone RPA-2.10 (300236, BioLegend, 1:200), IgD-AF488 clone 11-26c.2a (405718, BioLegend, 1:500), IgMb-BV510 clone AF6-78 (742344, BD OptiBuild, 1:500), TACI-AF647 clone 8F10 (558453, BD Biosciences, 1:500), CD45.2-BV786 clone 104 (563686, BD Horizon, 1:500), CD19-AF700 clone 1D3 (557958, BD Pharmingen, 1:500).

### Quantum Dot coupling of antibody

Quantum Dot (QD) antibody coupling was done using SiteClick Qdot 655 Antibody Labeling Kit (Molecular Probes, S10453) according to manufactures instructions. In brief, antibody (either “14D12” rat IgG2a to mouse MBL-C (Hycult Biotech), or “RTK2758” rat IgG2a isotype control (Abcam)) was concentrated in antibody preparation buffer to a concentration of 2 mg/mL or above. Next, carbohydrates on the antibody were modified by the incubation with β-galactosidase for 4 h at 37°C. Azide modification was achieved through incubation with uridine diphosphate glucose-GalT enzyme overnight at 30°C. Antibody with modified carbohydrates was purified and concentrated through a series of centrifugation steps using a molecular-weight cutoff membrane concentrator, and the buffer was simultaneously changed to 20 mM Tris, pH 7.0. Finally, 5ʹ-dibenzocyclooctyne-modified QD nanocrystals were coupled overnight at 25°C and stored at 4°C until further use.

### Nanoparticle Tracking Analysis

Samples for Nanoparticle Tracking Analysis (NTA) were analyzed using a NanoSight NS300 system (Malvern Panalytical) as previously described (Juul-Madsen et al., 2021). The system was configured with a 405 nm laser, a high-sensitivity scientific complementary metal–oxide– semiconductor Orca Flash 2.8/Hamamatsu C11440 camera (Malvern Panalytical), a syringe pump, and for fluorescence measurements, a 650 nm long-pass filter was used. The sample chamber was washed twice with 1 mL PBS with 1 mM EDTA (PBS/EDTA) before each measurement. All samples were thoroughly mixed before measurement and were injected into the sample chamber using 1-mL syringes. The measurement script comprised temperature control at 23°C, followed by a 20 s flush at a flowrate mark 1000. Next, sample advancement was stabilized by a 120 s advancement at flowrate mark 10. Recordings were captured continuously during a steady flow at flowrate mark 10 with five 60-s recordings separated by 5-s lag time between each sample. The videos were collected and analyzed using NanoSight software (version 3.4 with a concentration upgrade; Malvern). Automatic settings were used for the minimal expected particle size, minimum track length, and blur setting. Camera sensitivity and detection threshold were adjusted according to sample composition and kept constant for all samples to be directly compared. For fluorescence mode, the camera level was set to maximum (mark 16), and the detection threshold was set to minimum (mark 2). Serum samples from mice were analyzed in a 1:20 dilution in PBS/EDTA with a 1:20,000 dilution of MBL-C–specific (14D12) or isotype Antibody-QD reporters. A 50 nm cutoff was established for all samples to exclude unbound QD conjugates as well as QD conjugates bound to smaller forms of MBL-C.

### Immunohistochemical labelling of kidney tissue

After perfusion fixation, kidneys were stored in PBS with 30% sucrose and 0.1% sodium azide. For paraffin-embedding, the kidneys were washed in 10 mM PBS several times, dehydrated in 70%, 96%, and 99% ethanol for 2 hours, respectively, before they were transferred to xylene overnight. The day after, kidneys were embedded in paraffin. Paraffin embedded kidney sections (2 µm) were cut on a Leica RM 2165 microtome (Leica, Wetzlar, Germany) and dried at 60°C for 1 h. For immunofluorescent (IF) labelling, sections were placed in xylene overnight, rehydrated in graded alcohols, and heated in TEG buffer (10 mM Tris, 0.5 mM EGTA buffer, pH 9) at ∼100°C for 10 min to induce epitope retrieval. Sections were subsequently cooled for 30 min, incubated in 50 mM NH_4_Cl in 0.01 mM PBS for 30 min, and incubated at 4°C with primary antibody overnight. The following day, sections were washed, incubated with secondary antibody for 1 h, and coverslips were mounted using mounting medium (Dako fluorescence Mounting medium, #S3023). Immunofluorescent images were acquired by a confocal laser-scanning microscope (LSM 800 with Airyscan, Carl Zeiss Gmbh, Jena, Germany) and processed using ZEN lite 3.4 (Blue edition). For immunoperoxidase (IP) labelling, sections were prepared as stated above. In addition, endogenous peroxidase was blocked by placing sections in 30% H_2_O_2_ in methanol for 30 min and peroxidase-conjugated secondary antibodies were used. Furthermore, the sections were incubated with DAB, counterstained with Mayer’s haematoxylin (Sigma-Aldrich), dehydrated in graded alcohols, cleared in xylene and mounted with coverslips using Eukitt (Sigma-Aldrich). For periodic acid-Schiff (PAS) stainings, sections were incubated in 1% periodic acid for 10 min and treated with Schiff’s reagent for 20 min. Subsequently, all sections were counterstained with Mayer’s hematoxylin for 5 min and mounted with coverslips using Eukitt. Peroxidase-labelled and PAS-stained images were collected by a Leica DFC320 camera (Leica, Wetzlar, Germany). H&E and PAS stained kidney sections were scored by a histopathologist, blinded to the identity of the samples. High-resolution images of kidney and spleen sections were obtained using a Zeiss LSM 800 Airyscan confocal microscope with 4 lasers (405 nm, 488 nm, 561 nm, 640 nm). ZEN software was used for quantification of immunofluorescence kidney stainings. The following primary antibodies were used: anti-C3 (ab11887, Abcam, 1:25 (IP), 1:50 (IF)), anti-IgG2c (1079-08, SouthernBiotech, 1:25 (IF), 1:50 (IP)), anti-Nephrin (GP-N2, Progen, 1:100), anti-Ig (1010-08, SouthernBiotech, 1:100 (IF), 1:200 (IP)). The following secondary antibodies were used: Donkey-a-rabbit AF488 (A21206, ThermoFisher, 1:300), Streptavidin AF647 (405237, BioLegend, 1:500), Goat-a-Guinea Pig AF488 (A11073, ThermoFisher, 1:300), Goat-a-rabbit HRP (P0448, Dako, 1:200), Streptavidin HRP (P0397, Dako, 1:200)

### Time-resolved immunofluorometric analysis (TRIFMA) anti-dsDNA measurements

A FluoroNunc Maxisorp 96-well plate was coated with 100 μg/mL salmon sperm dsDNA (AM9680, Invitrogen) in PBS and incubated overnight at 4°C. Wells were blocked with 200 μL TBS containing 1% bovine serum albumin (BSA) (A4503, Sigma-Aldrich) for 1 hour at RT and washed 3 times with TBS/Tw (TBS containing 0.05% v/v Tween-20 (8.17072.1000, Merck)). Samples, standards and quality controls were diluted in TBS/Tw containing 5 mM EDTA and 0.1% w/v BSA, and were subsequently loaded onto the plate in duplicates. The plate incubated at 37°C for 1 hour. Then, wells were washed 3 times in TBS/Tw and incubated with biotinylated antibody (Table 2.5) at 37°C for 1 hour. Wells were washed 3 times in TBS/Tw, and Eu^3+^-tagged streptavidin (1244-360, PerkinElmer) diluted 1:1,000 in TBS/Tw containing 25 μM EDTA were subsequently added to the wells and incubated at RT for 1 hour. Finally, the wells were washed 3 times in TBS/Tw, 200 μL enhancement buffer (AMPQ99800, Amplicon) was added. The plate was shaken for 5 minutes and counts were read by a time-resolved fluorometry plate reader Victor X5 (Perkin Elmer).

### TRIMA Ig measurements

A FluoroNunc Maxisorp 96-well plate was coated with 1 μg/mL goat anti-mouse Ig in PBS and incubated overnight at 4°C. Wells were blocked in 1 mg HSA/mL TBS for 1 hour at RT and washed 3 times with TBS/Tw. Samples, standards and quality controls, diluted in TBS/Tw containing 100 μg/mL heat-aggregated human Ig, were subsequently loaded onto the plate in duplicates, and incubated overnight at 4°C. The wells were washed 3 times with TBS/Tw, and 1 μg/mL biotinylated goat anti-mouse Ig was added to the wells and incubated for 2 hours at RT. Wells were washed 3 times in TBS/Tw, and Eu^3+^-tagged streptavidin (1244-360, PerkinElmer) diluted 1:1,000 in TBS/Tw containing 25 μM EDTA were subsequently added to the wells and incubated at RT for 1 hour. Finally, the wells were washed 3 times in TBS/Tw, 200 μL enhancementbuffer (AMPQ99800, Amplicon) was added. The plate was shaken for 5 minutes and counts were read by a Victor X5 time-resolved fluorometry plate reader (Perkin Elmer).

### Immunofluorescence staining of spleens and auricular lymph nodes

A Cryostar NX70 Cryostat (ThermoFisher) was used to cut 16 μm thick spleen sections or 20 μm thick auricular lymph node sections which were mounted on SuperFrost+ glass slides (Fisher Scientific). Spleen sections were either acetone or PFA fixed, auricular lymph nodes were PFA fixed. For acetone fixation, the spleen samples were rinsed in PBS and fixed in acetone for 10 minutes at room temperature (RT), whereafter the slides were rehydrated in PBS for 3 minutes. For PFA fixation protocols, the slides were washed in PBS, fixed with 4% w/v PFA for 30 min at RT, incubated in TBS (10 mM Tris, 140 mM NaCl, pH 7.4) for 30 min at RT, rinsed briefly with PBS, and incubated with permeabilization buffer (PBS, containing 2% v/v FBS, 0.1% w/v sodium azide, 0.1% v/v Triton-X100) for 45 minutes at RT. Antibodies were diluted in staining buffer (PBS, 2% v/v FBS, 0.1% w/v sodium azide). The antibody mix was centrifuged at 10,000 *g* for 5 minutes and added onto the spleen samples, where it incubated overnight at 4°C. The slides were washed once with staining buffer for 5 minutes and washed 3 times in PBS with 0.01% v/v Tween-20 for 5 minutes. Slides were spot-dried and mounted using Fluorescence Mounting Medium (S3023, Dako). Imaging for quantification of GC formation was performed using an Olympus VS120 Upright Widefield fluorescence slide scanner equipped with a digital monochrome camera (Hamatsu ORCA Flash4.0V2) and a 2/3” CCD camera, as well as single-band exciters and a filter wheel with single-band emitters (Hoechst, FITC, Cy3, Cy5, and Cy7). Fiji v. 2.1.0/1.53c was used for image processing. The following antibodies were used: CD45.1-FITC clone A20 (110706, BioLegend, 1:300), CD45.2-APC clone 104 (109814, BioLegend, 1:300), CD138-PE clone 281-2 (142504, BioLegend, 1:500), CD169-PE clone 3D6.112 (142404, BioLegend, 1:500), IgD-AF488 clone 11-26c.2a (405718, BioLegend, 1:500), Ki67-eflour660 clone SolA15 (50-5698-82, Thermo Fisher Scientific, 1:500), CD21/35-PB clone 7E9 (123414, BioLegend, 1:500). Channel intensities were adjusted for visual clarity in represented micrographs, but quantification was performed on raw images throughout.

### Purification of B cells

The spleen was harvested, stored in MACS buffer (PBS, 2% FBS, 2 mM EDTA), the cells were mechanically dissociated and filtered through 70 μm cell strainers, whereafter the cell suspension was topped up with MACS buffer until 25 mL and filtered through 70 μm cell strainers again. The cell suspension was centrifuged at 200 *g* for 10 minutes at 4°C, resuspended in 5 mL RBC lysis buffer (155 mM NH_4_Cl, 12 mM NaHCO_3_, 0.1 mM EDTA), and incubated for 3 minutes at RT. The reaction was stopped by adding 47 mL MACS buffer, and centrifuged at 200 *g* for 5 minutes at 4°C. The supernatant was discarded and resuspended in 3 mL MACS buffer. B cell kit (Miltenyi Biotec, 130-090-862) was followed according to the manufacturer’s protocol.

### iGB cultures

NB21 feeder cells, kindly provided by Garnett Kelsoe, Duke University (Kuraoka et al., 2016), were seeded into 6-well plates at a density of 520 cells/cm^2^ for IL-4 stimulated wells and 5200 cells/cm^2^ for the four other conditions. The following day (day 0), B cells were purified according to the described protocol, and resuspended in B cell medium (BCM) (RPMI-1640 supplemented with 10% FCS, 55 μM 2-ME, 1% Pen/Strep, 1% MEM NEAA, 10 mM HEPES, 1 mM Sodium Pyruvate). B cells were pre-diluted in BCM with the given cytokine cocktail as indicated in figure legends. From day 2-8, 2/3 of the total volume of BCM was collected and fresh BCM was added to reach the same final volume. One mL of medium from each well of the 6 well plates was collected on the final day (day 10) for TRIFMA analyses. Cells were analyzed using flow cytometry, as described above. IL-4 (214-14, PeproTech) were used for stimulation

### Statistical analyses

GraphPad Prism v. 8.4.3 was used for statistical analyses. For each dataset, both tests for normality (such as Shapiro Wilk’s test) and Q-Q plots were used to determine whether the data were normally distributed. Data that were not normally distributed were log-transformed and re-tested for normality. The following data sets were log-normally distributed: Fig 1B, 1D, 1E, 1N, 1P, 3F, 5C, 5F, 5G, 5H, 6H, 6L-O, S6L-O. For Fig 6B, 6G, S6D-S6E, and S6H, Mann-Whitney test was used to analyze the data that were not normally distributed. All other datasets were normally distributed, except for the data in Figure 1L, 5F, and S6K which were neither normally, nor log-normally distributed. However, a non-parametric t-test for the isolated data between each group in panel 1L, 5F and S6K showed similar results (for Fig 1L: a significant increase in GC formation in Cre− between Unt and R848. For Fig 5F: a significant increase in CD8 frequencies in the IngLNs but not in other tissues. For S6K: slight, but significant increases across spleen, IngLN and blood, with a trend towards an increase in MesLN when tissues are analyzed individually), indicating that the observed differences, beyond being biologically robust and in agreement with the flow data, were also statistically robust. Parametric tests were used in all analyses, with specific tests indicated in the figure legends. All data is presented as bar graphs with mean ± SD. A p-value <0.05 was considered to be statistically significant. ns = p≥0.05, * = p<0.05, **= p<0.01, *** = p<0.001.

**Figure 1.**
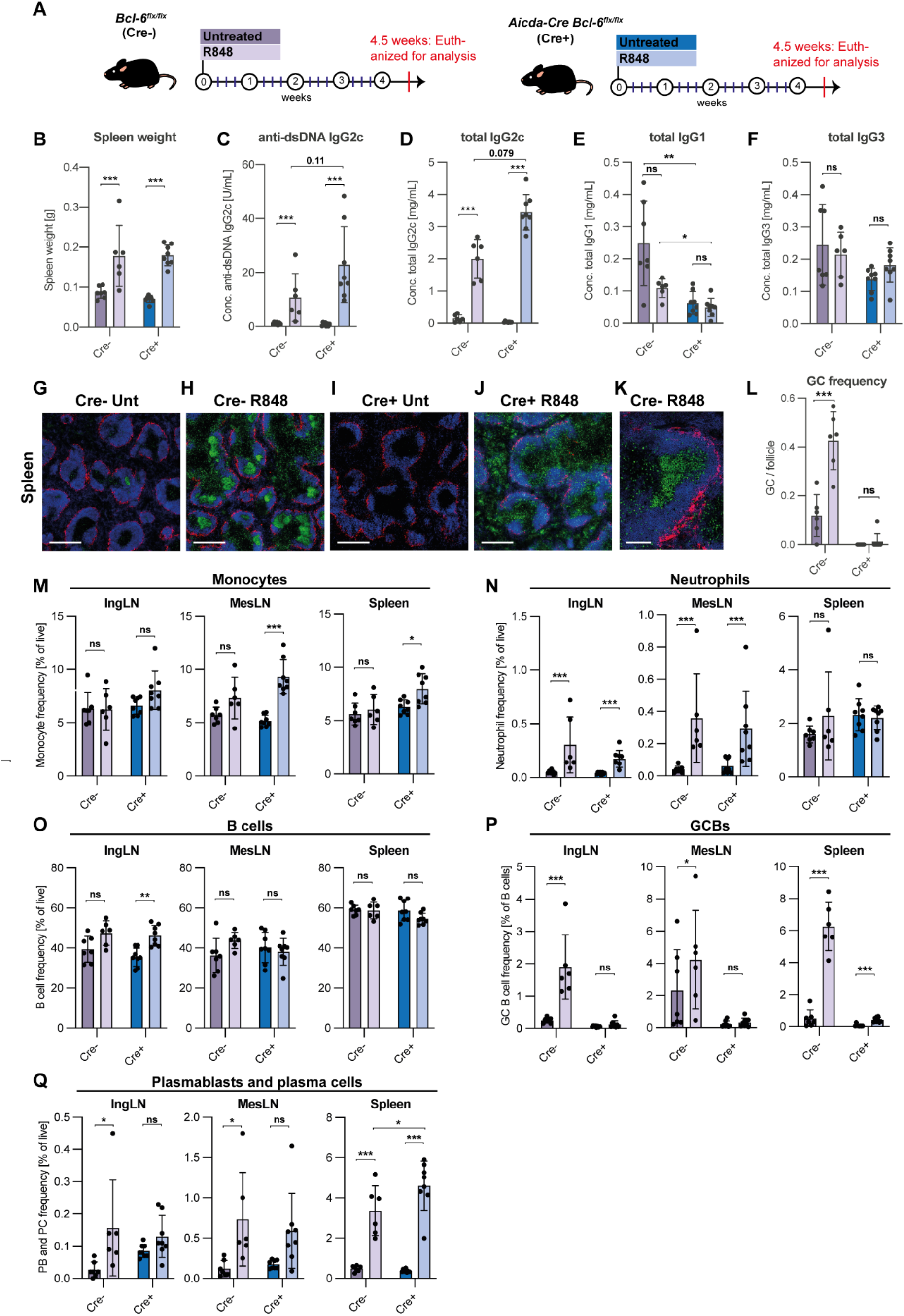
GC block causes increased levels of autoantibodies and PB/PCs in SLE-like mice. (**A**) Schematic overview of experimental setup and treatment protocol. Bcl-6^flx/flx^ (Cre−, purple) and Aicda-Cre Bcl-6^flx/flx^ (Cre+, blue) mice were either left untreated (dark color, n = 7 and n = 8, respectively) or treated with R848 (light color, n = 6 and n= 8, respectively), as indicated. (**B**) Spleen weights. (**C**) Anti-dsDNA IgG2c levels. (**D**) Total IgG2c levels. (**E**) Total IgG1 levels. (**F**) Total IgG3 levels. (**G**) Representative confocal micrograph of spleen from a Cre− untreated animal, stained for CD169 (red), IgD (blue) and Ki67 (green). Scale bar is 400 µm (**H**). As G, but for a Cre− R848-treated animal. (**I**) As G, but for a Cre+ untreated animal. (**J**) As G, but for a Cre+ R848-treated animal. (**K**) High-resolution image of a GC from a Cre− R848-treated mouse. Scale bar is 100 µm. **L**) GC per follicle in spleen. (**M**) Flow cytometry analyses of monocyte frequencies in IngLN, MesLN, and spleen (Ly6CG^int^ of live, singlet leukocytes). (**N**) As M, but neutrophil frequencies (Ly6CG^hi^ of live, singlet leukocytes). (**O**) As M, but B cell frequencies (B220^+^ CD4^-^ CD8^-^ of live, singlet leukocytes). (**P**) As M, but GCB frequencies (CD95^hi^ CD38^lo^ of B cells). (**Q**) As M, but PB and PC frequencies (CD138^hi^ of live, singlet leukocytes). Data are pooled from two independent experiments. Bar graphs show mean ± SD. Two-way ANOVA with Holm-Sidak’s post hoc test was used to analyze the data. ns = p≥0.05, * = p<0.05, ** = p<0.01, *** = p<0.001.

## Results

### Genetic GC block exacerbates autoimmune progression in a lupus model

Chronic epicutaneous application of the synthetic, small-molecule TLR7 agonist, R848 (Resiquimod), has previously been demonstrated to induce a robust lupus-like autoimmune phenotype in multiple genetic background strains of *Mus musculus* (Yokogawa et al., 2014). Leveraging this model, we set out to investigate the relative importance of the extrafollicular and GC pathways in the autoimmune response. To this end, we employed a transgenic Cre driver line displaying expression of Cre under the Aicda promotor (Kwon et al., 2008). We combined this with a conditional Bcl-6 knock-out line (Hollister et al., 2013), to achieve deletion of Bcl-6 specifically in GC B cells. As Bcl-6 is a master transcriptional regulator of the GC fates across GC B cells, T_FH_ cells and T_FR_ cells, this approach specifically prevents GC B cell differentiation, without affecting Bcl-6 dependent T cell subsets essential to support both GC and extrafollicular responses (Lee et al., 2011). As controls, we employed Cre negative Bcl-6^flx/flx^ littermates. In prior studies, when mice were treated with R848 3 times per week for 8 weeks, they displayed severe kidney damage, and 12 weeks of treatment led to a dramatic degree of mortality (Yokogawa et al., 2014). As we wanted to investigate the importance of pathways in production of autoantibodies and cell frequencies in the absence of secondary effects caused by organ failure, we decided to treat mice 3 times weekly for only 4 weeks (**Fig. 1A**).

After 4 weeks of treatment, the Bcl-6^flx/flx^ (Cre−) controls and the Aicda-Cre Bcl-6^flx/flx^ (Cre+) mice showed similar, significant increases in spleen weight upon R848 treatment (**Fig. 1B**). Anti-dsDNA autoantibodies of the IgG2c subtype measured in serum were dramatically elevated upon R848 treatment compared with untreated animals. Surprisingly, there was a trend towards higher levels in treated Cre+ mice compared with treated Cre− mice (**Fig. 1C**). A similar trend towards an increase was seen in total IgG2c levels (**Fig. 1D**). No statistically significant differences in total IgG1 and total IgG3 levels were seen upon treatment (**Fig. 1E and F**).

To validate the effect of R848 treatment and the integrity of the GC block in Cre+ mice, we carried out immunofluorescence staining of spleens to identify GC formation (**Fig. 1G-J**). Using the proliferation marker Ki67 and the naïve B cell marker IgD, it was evident that larger and more frequent GCs were observed upon R848 treatment in Cre− animals (**Fig. 1G, H and K**). Quantification revealed that this difference was statistically significant (**Fig. 1L**). Cre+ animals did not display any baseline or R848-induced GCs (**Fig. 1I and J**). However, in Cre+ treated mice we did observe many proliferating cells at the T-B border and in the red pulp (**Fig. 1J**), likely abortive primary foci and extrafollicular foci, respectively.

Flow cytometric analyses of inguinal LNs (IngLNs), mesenteric LNs (MesLN), and the spleen were carried out (**Fig. 1M-Q, Fig S1A**). In treated animals, we saw slight increases in monocyte and neutrophil frequencies in some of the tissues (**Fig. 1M and N**), and a slight increase in B cell frequencies in the skin-draining IngLNs (**Fig. 1O**), which might be caused by the direct stimulatory effect from the R848 treatment of the ear skin. We observed robust GC B cell frequencies in Cre− R848 treated mice, compared to untreated littermates (**Fig. 1P**). No GC B cells were found in Cre+ animals, further validating the fidelity of the GC block (**Fig. 1P**). Surprisingly, despite this, we found a significant increase in PB/PC frequencies upon treatment in both groups, and the level was significantly higher in the spleens of mice harboring a GC block compared to Cre− R848-treated littermate controls (**Fig. 1Q**). Splitting up for PB and PC, it was clear that this difference between the two groups for the spleen was attributable to PB (**Fig. S1B**) rather than PC (**Fig. S1C**) expansion in Cre+ animals, as compared to Cre− littermates. This observation corresponded well with the increase in plasma IgG2c autoantibody levels as well as total IgG2c levels (**Fig 1C and D**). Taken together, this surprisingly indicated an exacerbated autoimmune phenotype in GC block mice compared to GC sufficient mice upon R848 treatment.

**Supplementary Figure 1.**
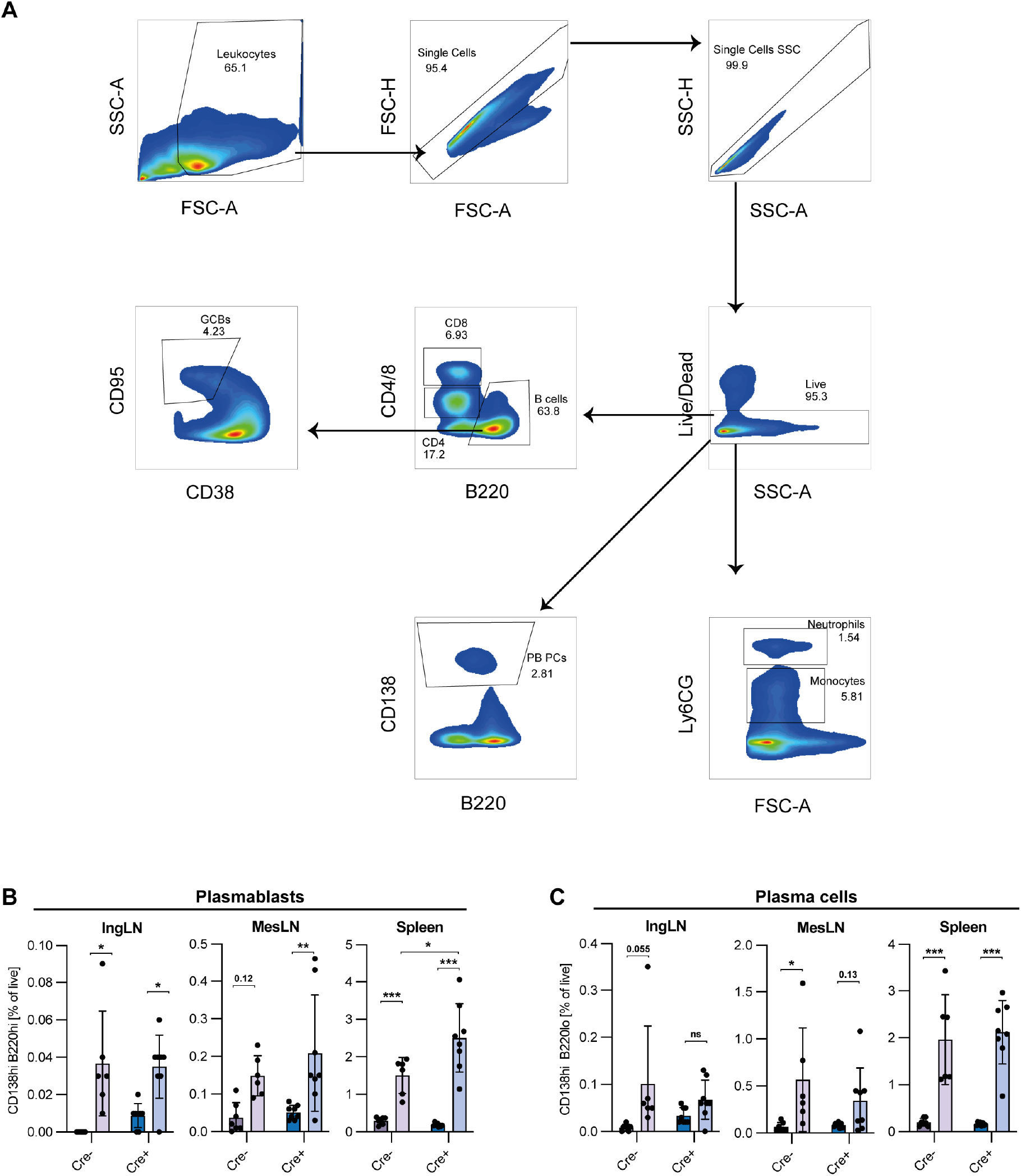
Gating strategy for R848 cohorts, PB and PC frequencies. (**A**) Leukocytes were gated based on size (FSC-A) and granularity (SSC-A). Doublets were excluded with two singlet gates, first FSC-H vs. FSC-A, and then SSC-H vs. SSC-A. After that, dead cells were excluded. From the live gate, monocytes and neutrophils were gated using the Ly6C/G marker. Moreover, B cells were selected, from which GC B cells were selected based on CD95 expression and the absence of CD38 expression. PB and PCs were selected from the live gate based on CD138. (**B**) PB (CD138^hi^ B220^hi^ of live, singlet leukocytes) frequencies across secondary lymphoid tissues. (**C**) PC (CD138^hi^ B220^lo^ of live, singlet leukocytes) frequencies across secondary lymphoid tissues. Data in B and C are pooled from two independent experiments. Bar graphs show mean ± SD. Two-way ANOVA with Holm-Sidak’s post hoc test was used to analyze the data. ns = p≥0.05, * = p<0.05, ** = p<0.01, *** = p<0.001.

To further understand the local effects of R848 treatment, we performed immunofluorescence microscopy analyses of draining auricular lymph nodes (AurLNs) from treated mice and untreated controls. We observed gross enlargement of the lymph nodes of treated animals, with a robust induction of GCs in Cre− R848-treated mice (**Fig. S2A**). In comparison, the Cre+ R848-treated mice had many proliferating cells outside the follicles (**Fig. S2A**). These dividing cells in the AurLNs overlapped to some extent with the PC marker CD138, pointing towards dividing extrafollicular PCs (**Fig. S2B**).

**Supplementary Figure 2.**
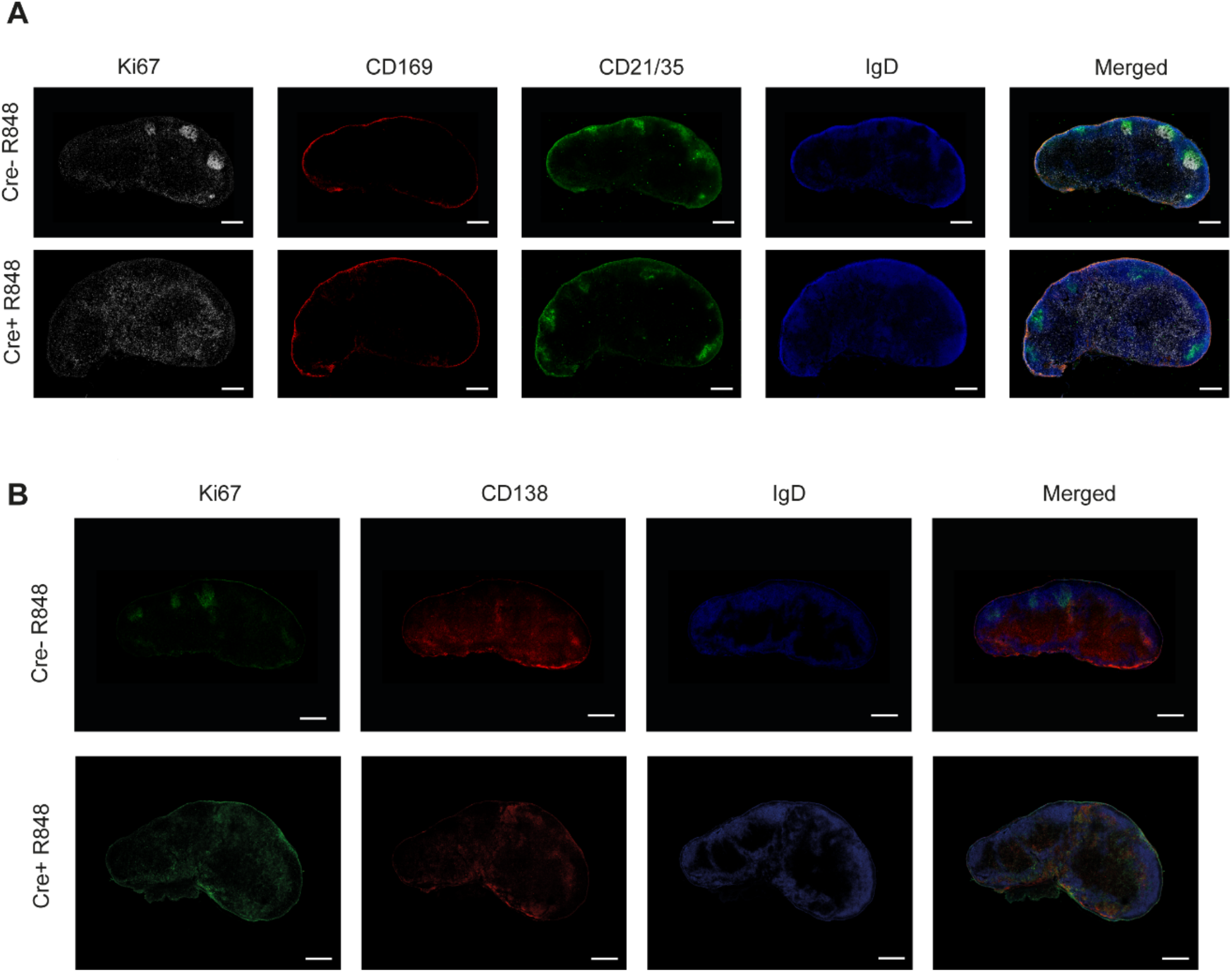
Immunofluorescence stainings show dividing cells in the paracortex of the AurLN which overlaps with CD138 staining. (**A**) The following targets were stained for: Ki67 (white), CD169 (red), CD21/35 (green), IgD (blue). (**B**) The following targets were stained for: Ki67 (green), CD138 (red), IgD (blue). Scale bar is 350 μm. Images were generated by tile scanning with a 10% overlap using a mechanical stage, followed by stitching. Channel intensities have been adjusted for visual clarity.

### Nanoparticle tracking analyses

Using nanoparticle tracking analyses, we recently identified unique superoligomeric complexes (spMBL) formed between cell-free DNA and mannan-binding lectin (MBL), as a hallmark in blood samples from SLE patients and lupus mice (Juul-Madsen et al., 2021). These spMBL complexes correlated with disease activity in SLE patients, and correlated with formation of GCs and drove loss of immunological tolerance in a murine lupus model (564Igi). To further understand the importance of the increased anti-dsDNA IgG2c autoantibodies in Cre+ R848-treated mice, in the face of a complete absence of GCs, we implemented this nanoparticle tracking approach (**Fig. 2A-C**). We analyzed superoligomeric complexes in the band from 90-130 nm (**Fig. 2D and E**). In agreement with spMBL as a lupus marker, treated mice tended towards higher levels, as compared to untreated mice, but interestingly, we also observed a global trend towards higher levels in Cre+ compared to Cre− animals (**Fig. 2F**). Although they did not reach statistical significance, these observations were well in line with the previously noted increases in anti-dsDNA and total IgG2c antibodies upon R848 treatment, and in Cre+ compared to Cre− animals (**Fig. 1C**).

**Figure 2.**
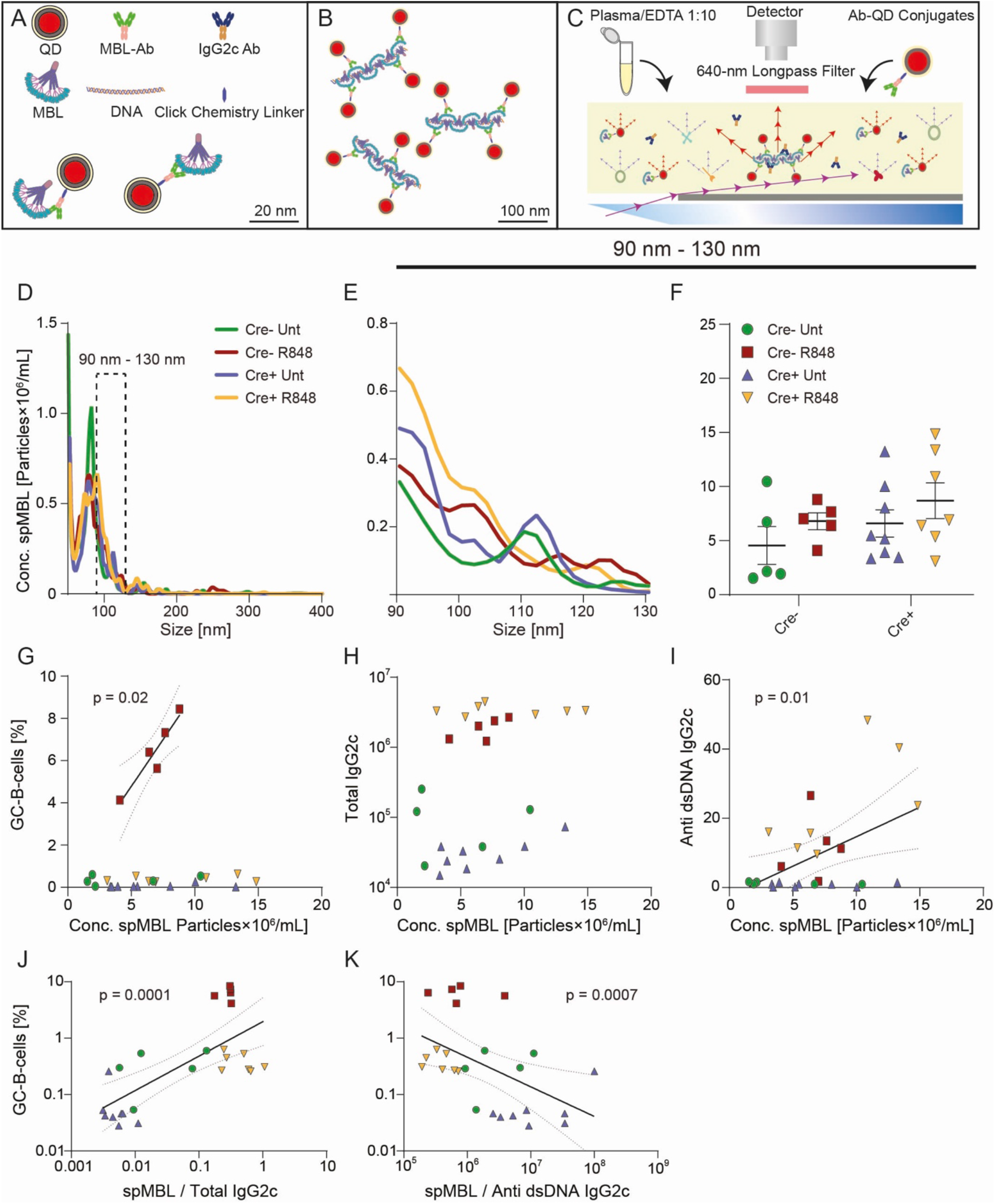
Trend towards increased levels of spMBL particles in serum from Cre+ R848 treated mice, and GC and autoantibody correlations with spMBL. (**A-C**) Schematic overview of experimental setup for spMBL analysis of serum samples. (**D**) Samples were tested for the size interval 90-130 nm from Aicda-Cre Bcl-6^flx/flx^ (Untreated: blue, R848-treated: yellow) and Bcl-6^flx/flx^ littermate controls (Untreated: green, R848-treated: red). (**E**) Zoom onto the range of 90 to 130 nm. (**F**) Graph showing mean ± SEM and individual measurements across treatment protocols for Cre− untreated (n= 5), Cre− R848-treated (n=5), Cre+ untreated (n=8), and Cre+ R848-treated (n=7) mice. (**G**) Correlation analysis of GCB vs. Conc. spMBL. (**H**) Correlation analysis of total IgG2c levels vs. Conc. spMBL. (**I**) Correlation analysis of anti-dsDNA IgG2c vs. Conc. spMBL. (**J**) Correlation analysis of GCB cells vs. spMBL/total IgG2c (**K**) Correlation analysis of GCB vs. spMBL/anti-dsDNA IgG2c. Two-way ANOVA with Holm-Sidak’s post hoc test was used to analyze the data in F. Linear regression models were used to analyze G-K. ns = p≥0.05, * = p<0.05, ** = p<0.01, *** = p<0.001.

In our prior study on autoimmune mice carrying an autoreactive B cell receptor knock-in (564Igi) on a wild-type background (Juul-Madsen et al., 2021), a significant inverse correlation was found between the frequency of splenic GC B cells and the ratio between the spMBL and anti-dsDNA antibody concentrations measured in serum. This suggested that an excess of spMBL increased GC B cell formation while an excess of anti-dsDNA antibodies decreased GC B cell proliferation, or vice versa, indicating a potential negative feedback loop. In consideration of this, we next asked whether the same correlation could be established in the R848 model, and how this phenomenon was impacted in GC block mice.

In Cre− treated mice, GC B cell frequencies were positively correlated with the concentration of spMBL particles in serum (**Fig. 2G**). Total IgG2c levels were similar between treated Cre+ and Cre− mice (**Fig. 2H**) with a significant positive correlation between anti-dsDNA IgG2c and the level of spMBL particles (**Fig. 2I**). A significant positive correlation was also observed between the frequency of splenic GC B cells and the ratio between the spMBL and total IgG2c levels in serum (**Fig. 2J**). Conversely, a significant inverse correlation was found between the frequency of splenic GC B cells and the ratio between the spMBL and anti-dsDNA IgG2c antibody concentrations measured in serum (**Fig. 2K**). Taken together, these findings now took our original observations from the autoreactive B cell receptor knock-in model (564Igi) into the epicutaneous R848 model on wild-type (Cre−) background. Importantly, in Cre+ mice, there was an uncoupling of the concentration of spMBL particles and GC B cell levels, indicating that the GC block failed to curb the production of spMBL complexes.

### Enhanced immune complex deposition in kidneys of R848-treated mice

To understand the pathological importance of the GC block in R848-treated mice and the elevated serum IgG2c, serum anti-dsDNA IgG2c and spMBL levels, we investigated whether there were any pathological changes associated with lupus nephritis in the kidneys. First, we carried out Periodic acid-Schiff (PAS) staining, which displayed no obvious kidney injury upon R848 treatment (**Fig. 3A**). Apart from the presence of mesangial and capillary immune deposits, histopathological changes associated with lupus nephritis may include increased matrix or mesangial cellularity, endocapillary proliferation, thickening of capillary walls, glomerular tuft necrosis, extracapillary proliferation (crescents), karyorrhexis, hyaline thrombi (micronodular intracapillary aggregation of immune complexes), and glomerular sclerosis (segmental or global), as well as, rarely, pathognomonic hematoxylin bodies (Gasparotto et al., 2020; Weening et al., 2004). However, no significant histopathologic findings were identified by light microscopy in any of the mice included in any of the groups. In line with the PAS stainings, we did not observe any differences in glomerular nephrin levels among the groups, suggesting normal glomerular podocytes (**Fig. 3B**). As we could not identify any gross pathological changes nor changes in nephrin levels in the kidneys upon treatment, this verified our short-term treatment strategy in terms of the goal to investigate early immune-driven events in the absence of any secondary pathology.

**Figure 3.**
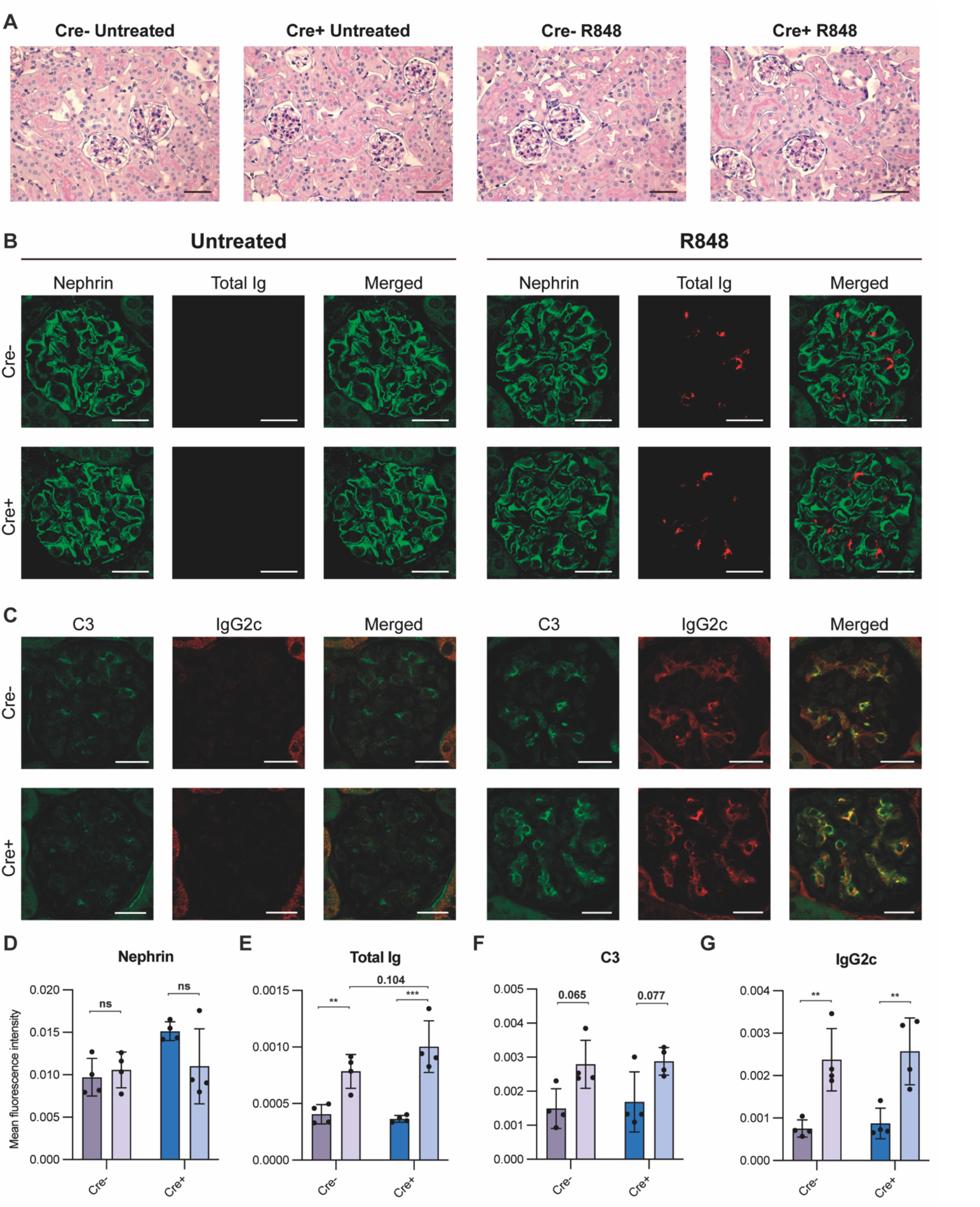
Kidney staining reveals immune complex deposition in R848-treated mice. (**A**) PAS stained kidney sections of Cre− untreated (n=4), Cre− R848-treated (n=4), Cre+ untreated (n=4), Cre+ R848-treated (n=4). Scale bar is 50 µm. (**B**) Immunofluorescence staining of kidney sections targeting nephrin (green) and total Ig (red). (**C**) Immunofluorescence staining of kidney sections targeting C3 (green) and IgG2c (red). (**D**) Quantification of immunofluorescence staining targeting nephrin, (**E**) total Ig, (**F**) C3, (**G**) and IgG2c. Scale bar is 20 µm. Two-way ANOVA with Holm-Sidak’s post hoc test was used to analyze the data. Bars show mean ± SD. ns = p>0.05, * = p < 0.05, **= p<0.01, *** = p < 0.001.

To evaluate if immune complex deposition occurred in the kidney glomeruli, and if there were any differences between GC-sufficient and deficient groups, we performed immunofluorescence staining of kidney sections targeting total Ig, C3, and IgG2c (**Fig 3B, 3C, 3E-G**). The total Ig levels in glomeruli were clearly increased upon R848-treatment. Interestingly, we found a trend towards an increase in the total Ig levels of Cre+ mice compared with littermate R848-treated controls (**Fig. 3B and E**). We also found that there was a significant increase of antibodies of the pathogenic subtype IgG2c upon R848 treatment (**Fig. 3C and G**), and a trend towards an increase in C3 deposition upon R848-treatment (**Fig. 3C and F**). However, differences between Cre+ and Cre− R848-treated groups were seen neither in C3 nor IgG2c levels (**Fig. 3F and G**).

Taken together, R848-treated mice displayed immune complex deposition in glomeruli, based on an increased level of C3, total Ig and IgG2c (**Fig. 3E-G**). We corroborated these findings by peroxidase-staining, as a corollary to the immunofluorescence microscopy, and this confirmed the glomerular changes in total Ig, C3 and IgG2c upon R848-treatment (**Fig S3**). Based on the immunofluorescence and immunoperoxidase findings of immune complex deposition (Ig, IgG2c and C3), in the absence of gross histopathological changes of the kidneys, this corresponds to Class I pathology (minimal mesangial lupus nephritis) displaying mesangial immune deposits without mesangial hypercellularity, as defined by the International Society of Nephrology (ISN) and the Renal Pathology Society (RPS) 2004 classification system (Weening et al., 2004).

**Supplementary Figure 3.**
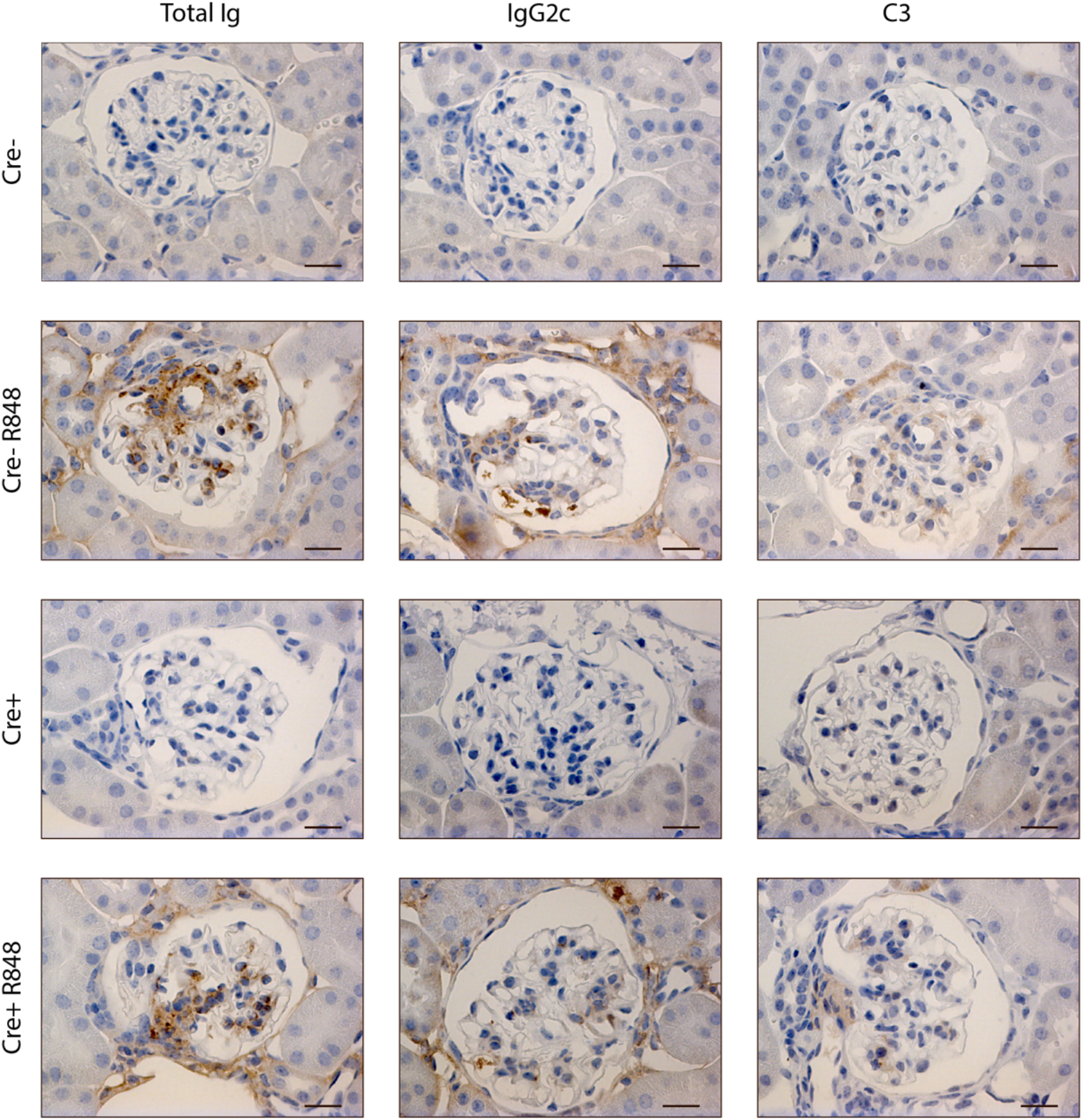
Increased immune deposits in glomeruli of R848-treated mice. Representative images of immunohistochemical peroxidase staining targeting total Ig, IgG2c and C3 on sections from Cre− untreated (n=4), Cre− R848-treated (n=4), Cre+ untreated (n=4), Cre+ R848-treated (n=4) mice. Scale bars = 20 μm.

### An intrinsic GC block drives B cell differentiation into terminally differentiated PCs

To investigate whether a B cell intrinsic block of the GC pathway affects their capacity to differentiate into PBs and PCs, we leveraged a modified *in vitro* setup for induced GC B cell (iGB) cultures (Kuraoka et al., 2016; Nojima et al., 2011) (**Fig. 4A**). Naïve B cells purified from Cre− and Cre+ Bcl-6^flx/flx^ mice by negative magnetic-activated cell sorting were seeded onto fibroblast feeder cells expressing CD40L, IL-21 and BAFF, and stimulated with IL-4 (**Fig. 4A**).

**Figure 4.**
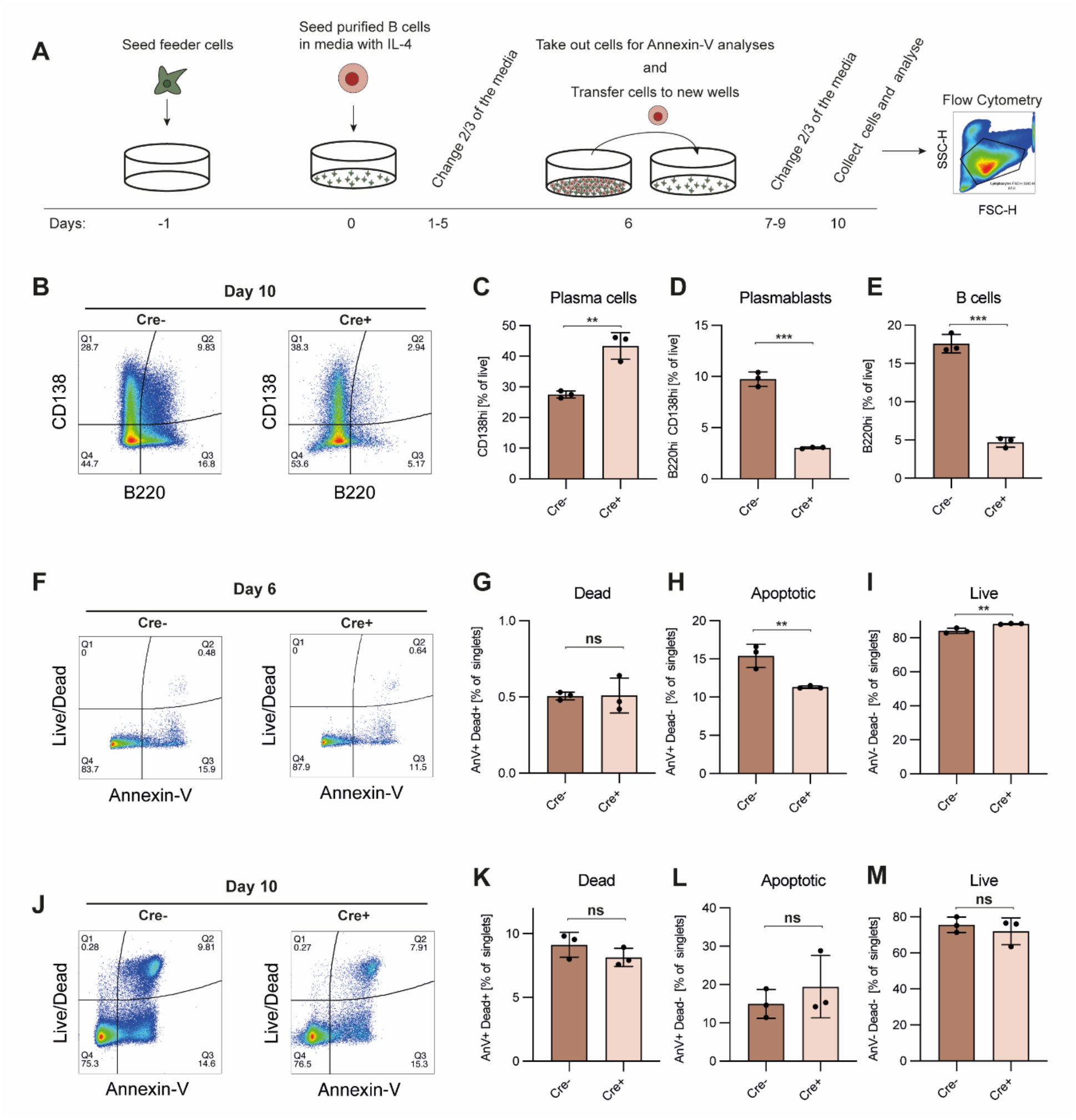
Bcl-6 deficient cells more readily differentiate into PCs in iGB cultures. B cells in iGB cultures were treated with IL-4 in conjunction with CD40L, BAFF, and IL-21. (**A**) Schematic overview of the iGB culture system. (**B**) Representative terminal CD138 vs. B220 bivariate plot for iGB cultured B cells stimulated with IL-4. (**C**) Bar graph showing B cell frequencies (B220+, CD138neg). (**D**) As C, but showing PB frequencies (B220+, CD138+). (**E**) As for C, but showing PC frequencies (B220neg, CD138+). Data are representative of three independent experiments with a cumulative 8 replicates in total. Bar graphs show mean ± SD. (**F**) Day 6 representative terminal Live/Dead vs. Annexin V bivariate plot for iGB cultured B cells stimulated with IL-4. (**G**) Bar graph showing dead cell frequencies (AnV-, Dead+). (**H**) As F, but showing apoptotic cell frequencies (AnV+, Dead-). (**I**) As for F, but showing live cell frequencies (AnV-, Dead-). (**J**) Day 10 representative terminal Live/Dead vs. Annexin V bivariate plot for iGB cultured B cells stimulated with IL-4. (**K**) Bar graph showing dead cell frequencies (AnV-, Dead+). (**L**) As for K, but showing apoptotic cell frequencies (AnV+, Dead-). (**M**) As for K, but showing live cell frequencies (AnV-, Dead-). Data for apoptosis assays are from one experiment with 3 replicates. Bar graphs show mean ± SD. Unpaired t-tests were used to analyze all datasets. ns = p≥0.05, * = p<0.05, ** = p<0.01, *** = p<0.001.

The combination of CD40L, BAFF, IL-21, and IL-4 stimulation has previously been shown to induce a robust expansion of B cells with a GC-like phenotype, followed by differentiation of the cells into PBs and finally PCs (Kuraoka et al., 2016; Nojima et al., 2011). Following stimulation of cultures with IL-4, we performed flow cytometric analyses and observed significantly higher B cell (B220+, CD138neg) frequencies in cultures derived from Cre− mice, compared to those derived from Cre+ mice (**Fig. 4B+E and Fig. S4A+B**). This was mirrored by a similar relative increase in PBs (B220+, CD138+) in Cre− cultures (**Fig. 4B and D**), but a relative decrease in PCs (B220neg, CD138+) (**Fig. 4B and C**). Of interest, the total number of cells in the live gate for Cre− cultures was approximately 4 times higher than that of Cre+ cultures (Cre−: 140,000 vs. Cre+: 33,000, **Fig. S4C**). To understand this difference in cell numbers, we investigated whether the Cre+, and hence Bcl-6 deficient, B cells had an increased propensity to undergo apoptosis, because Bcl-6 has previously been reported to suppress P53 and inhibit apoptosis in GC B cells (Phan and Dalla-Favera, 2004). Somewhat surprisingly, we found that upon IL-4 stimulation, there was no difference in the frequency of dead cells (**Fig. 4F and G**), a slight and significant drop in apoptotic cell frequency (**Fig. 4F and H**), and a corresponding increased relative frequency of live cells in Cre+ cultures compared to Cre− cultures on day 6 (**Fig. 4F and I**). However, at day 10 there were no significant differences in live, apoptotic, nor necrotic cell frequencies between Cre− and Cre+ cultures (**Fig. 4J-M**). Thus, apoptosis could not account for the dramatic difference in resulting cell numbers between Cre+ and Cre− (**Fig. S4C**). Taken together, this suggested that the higher overall cell numbers in Cre− cultures was not simply a reflection of increased apoptosis among Bcl-6 deficient cells in Cre+ cultures, but rather represented an improved intrinsic proliferative potential of the Bcl-6 sufficient cells.

In summary, our iGB experiments revealed a vigorous expansion of B cells and PCs in Cre− cultures, but less pronounced PC differentiation, whereas Cre+ cultures conversely displayed a lesser degree of proliferation but more pronounced PC differentiation. This suggested that B cells with an intrinsic GC block may differentiate quicker to PCs, and thereby lose their proliferative capacity, a notion that is in line with the established function of Bcl-6 in repressing upregulation of Blimp-1 (Vasanwala et al., 2002).

**Supplementary Figure 4.**
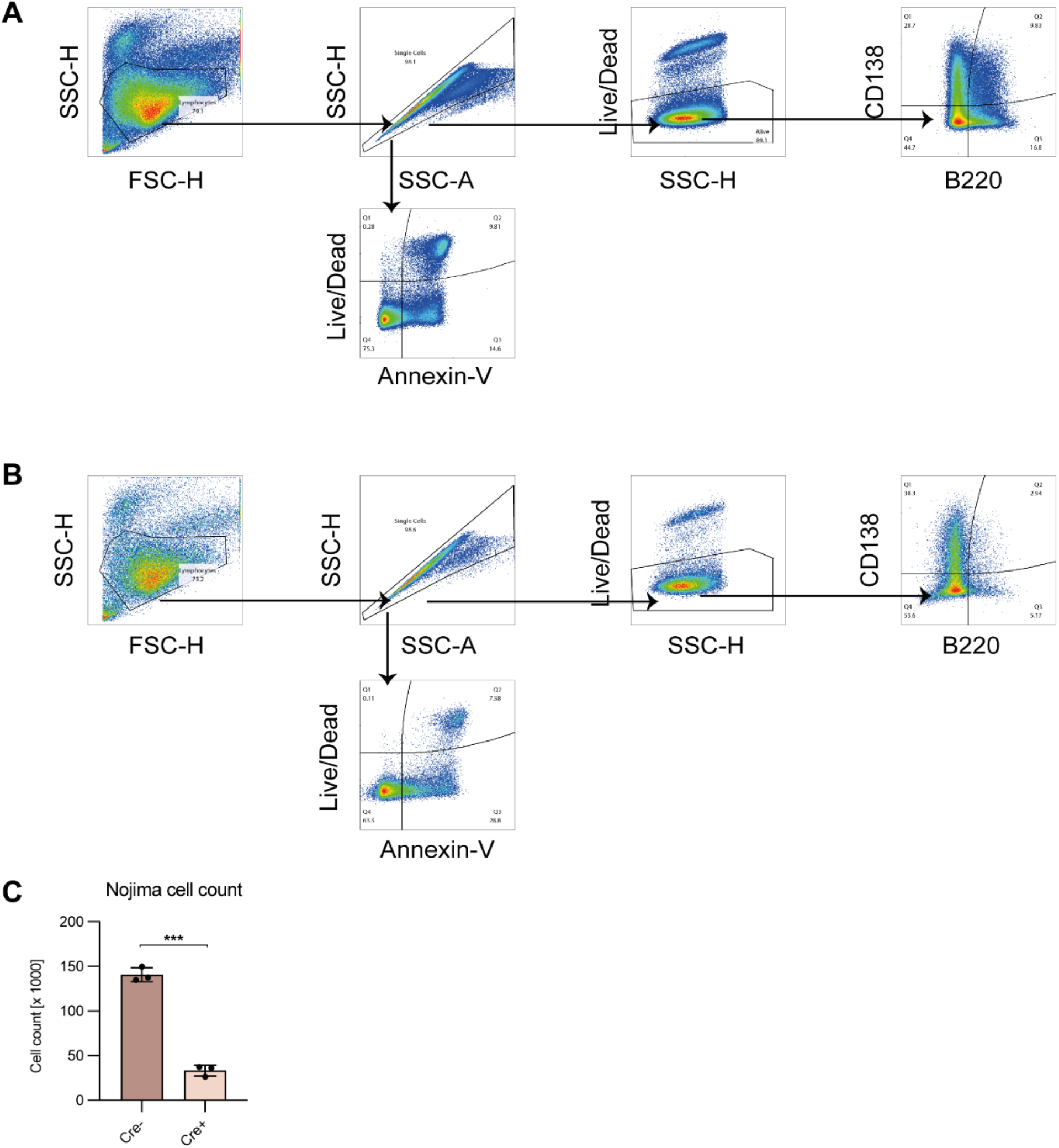
Gating strategy for Nojima cultures for (**A**) Cre−, (**B**) Cre+. Lymphocytes were gated based on size (FSC-A) and granularity (SSC-A). Doublets were excluded with a singlet gate (SSC-H vs. SSC-A). Apoptotic cells were gated based on the singlet gate using Annexin V and live/dead marker. Dead cells were excluded, and PCs, PBs and B cells were gated based on CD138 and B220 levels. (**C**) Live cell count from flow cytometry on day 10. Bar graph shows mean ± SD. ns = p≥0.05, * = p<0.05, **= p<0.01, *** = p<0.001.

### B cells harboring a GC block competitively contribute to the PC compartment in an autoreactive setting

Our observation that Bcl-6 deficient B cells rapidly lost their replicative potential *in vitro* (**Fig. 4**) was somewhat at odds with our *in vivo* observations from the R848 model, which displayed a global increase in PCs and autoantibodies (**Fig. 1**). This is because even if Bcl-6 deficient cells more readily became PCs, their poorer capacity to expand compared to Bcl-6 sufficient cells would be predicted to limit PC output. However, it was possible that the constant autoinflammatory drive in an *in vivo* setting where B cells were globally prevented from differentiating down the GC pathway would dysregulate the PC differentiation process and disproportionately drive extrafollicular differentiation. To address this possibility, we asked whether B cells with a GC block would be precluded from contributing to the PC pool in an environment where a large fraction of competitor B cells could form GCs. To achieve this, we leveraged a mixed bone marrow (BM) chimera model allowing interrogation of the competitive potential of B cells with a defined genetic defect in a lupus-like setting (Degn et al., 2017b; Wittenborn et al., 2021). In this model, an autoreactive B cell receptor knock-in clone (clone 564Igi) initiates an autoreactive process that subsequently recruits proto-autoreactive B cells from the non-564Igi B cell population. The B cells derived from the 564Igi compartment eventually are outcompeted and constitute only a minor fraction of the total B cell repertoire. Uniquely to this model, the spontaneous autoreactive GCs established by the 564Igi compartment become populated and chronically self-sustained by the non-564Igi (WT) B cells and gain independence from the initial 564Igi trigger. From around six weeks after reconstitution, GCs are almost exclusively (∼95%) composed of WT-derived cells (Degn et al., 2017b; Green et al., 2021). Reconstitution with a third of each of 564Igi BM, BM from a wild-type donor, and BM from a donor harboring a specified genetic defect, hence results in chimeras with two equal-sized compartments of B cells sufficient or deficient in the gene of interest. With the use of appropriate congenic markers, their competitive recruitment and participation in the autoreactive GC reaction and relative contribution to the PC compartment can subsequently be evaluated to elucidate the functional relevance of their intrinsic molecular differences. Accordingly, we set up mixed BM chimeras by irradiating WT CD45.1/1 recipients and reconstituting with 1/3 of each of 564Igi knock-in BM, WT CD45.1/1 BM, and either Aicda-Cre+ Bcl-6^flx/flx^ or Aicda-Cre− Bcl-6^flx/flx^ BM (**Fig. 5A and S5**).

**Figure 5.**
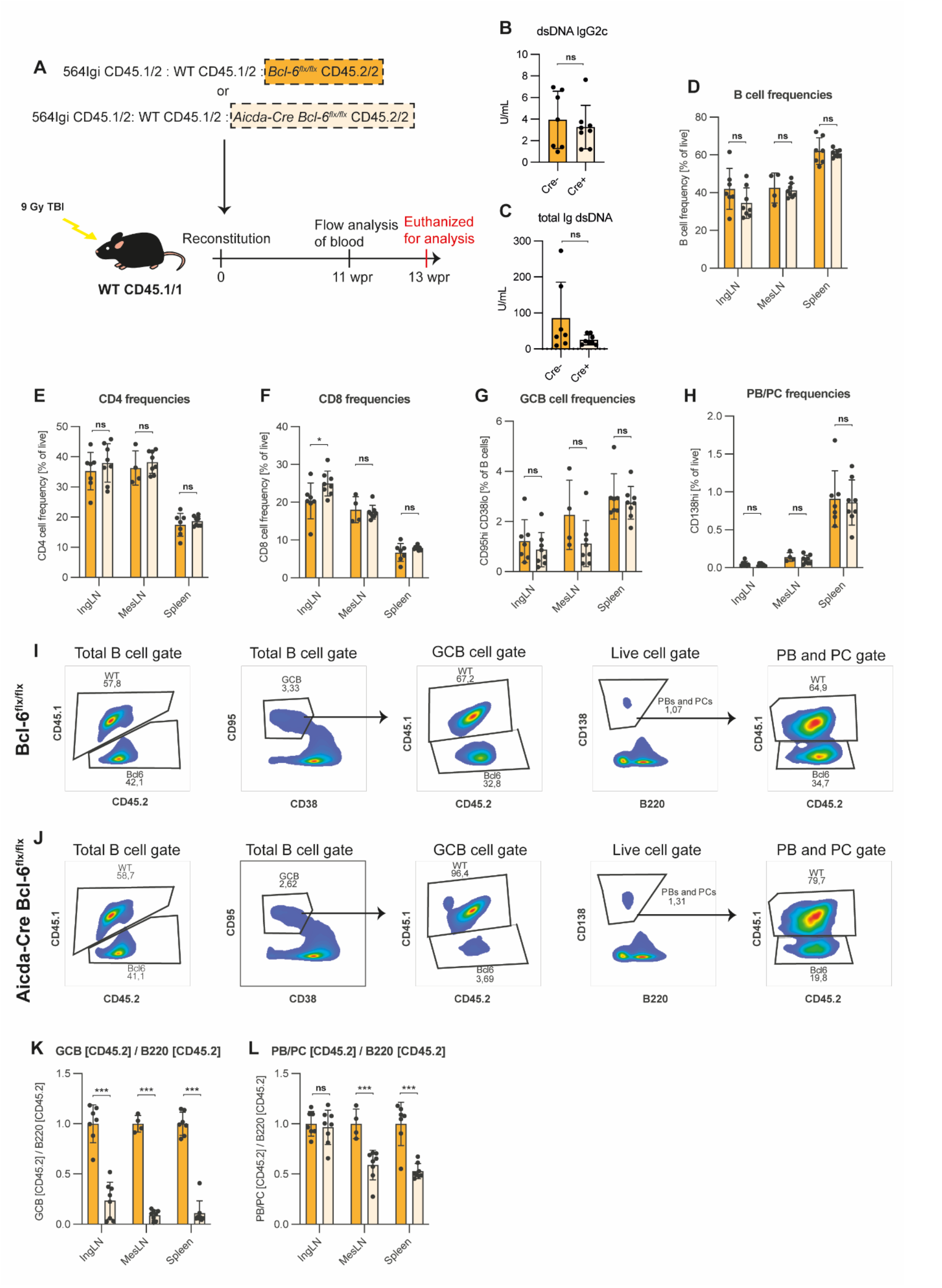
B cells harboring a GC block contribute to the PC lineage in a GC sufficient environment. (**A**) Schematic overview of the mixed bone marrow chimera setup. Lethally irradiated CD45.1/1 recipients were reconstituted with CD45.2/2 564Igi BM, WT CD45.1/2 BM and either Bcl-6^flx/flx^ (Cre−, orange, n = 7) or Aicda-Cre Bcl-6^flx/flx^ (Cre+, light orange, n=8). (**B**) dsDNA IgG2c TRIFMA. (**C**) total Ig TRIFMA. (**D**) Flow cytometric analysis of B cell frequencies (B220^+^ of live, singlets). (**E**) CD4 frequencies (CD4 of live, singlets). (**F**) CD8 frequencies. (**G**) GCB frequencies (CD95^hi^ CD38^lo^ of B cells). (**H**) PB/PC frequencies (CD138^hi^ of live, singlets). (**I**) Representative bivariate plots with gates for Bcl-6^flx/flx^ chimeras. (**J**) Representative bivariate plots with gates for Aicda-Cre Bcl-6^flx/flx^ chimeras. (**K**) Ratio of CD45.2+ of GCB to CD45.2+ of total B cells. (**L**) Ratio of CD45.2+ of PBs/PCs to CD45.2+ of total B cells. The results are obtained from a single experiment with the number of mice given above. Bar graphs show mean ± SD. ns = p≥0.05, * = p<0.05, **= p<0.01, *** = p<0.001.

There were no differences in the basic parameters when comparing Cre− control BM chimera mice with Cre+ BM chimera mice, in which approximately 50% of the B cells harbored a GC block (**Fig. 5B-H**). That is, aside from a very small but significant relative increase of CD8 T cells in IngLN of Cre+ chimeras, we saw no statistically significant differences in anti-dsDNA IgG2c (**Fig. 5B**), total anti-dsDNA Ig (**Fig. 5C**), B cell frequencies (**Fig. 5D**), CD4 and CD8 T cell frequencies (**Fig. 5E and F, respectively**), overall GC B cell frequencies (**Fig. 5G**) and PB/PC frequencies (**Fig. 5H**) between the groups. This confirmed that the two groups of chimeras were comparable and had robust GCB and PC compartments. In the total B cell compartment, CD45.2 positive cells were present at levels comparable to that of CD45.1 cells in both Cre− (**Fig. 5I**) and Cre+ (**Fig. 5J**) chimeras. However, within the GC B cell gate, CD45.2 cells were robustly represented in Cre− chimeras, but virtually absent in Cre+ chimeras (**Fig. 5I and J**). When quantifying this effect across chimeras and expressing as the ratio of CD45.2 of GCB relative to CD45.2 of total B cells, it was clear that Bcl-6 deficient cells, as expected from their genetic deficiency, were incapable of contributing to the GC compartment (**Fig. 5K**). However, when similarly comparing PB/PC ratio over B cells, the cells harboring a GC block remained able to contribute to the final PB/PC pool, albeit underrepresented relative to the competitor cells (**Fig. 5L**). Taken together, these findings demonstrated that in a GC-sufficient environment, B cells experiencing a block in their ability to partake in the GC reaction readily contributed to the PB and PC compartments.

**Supplementary Figure 5.**
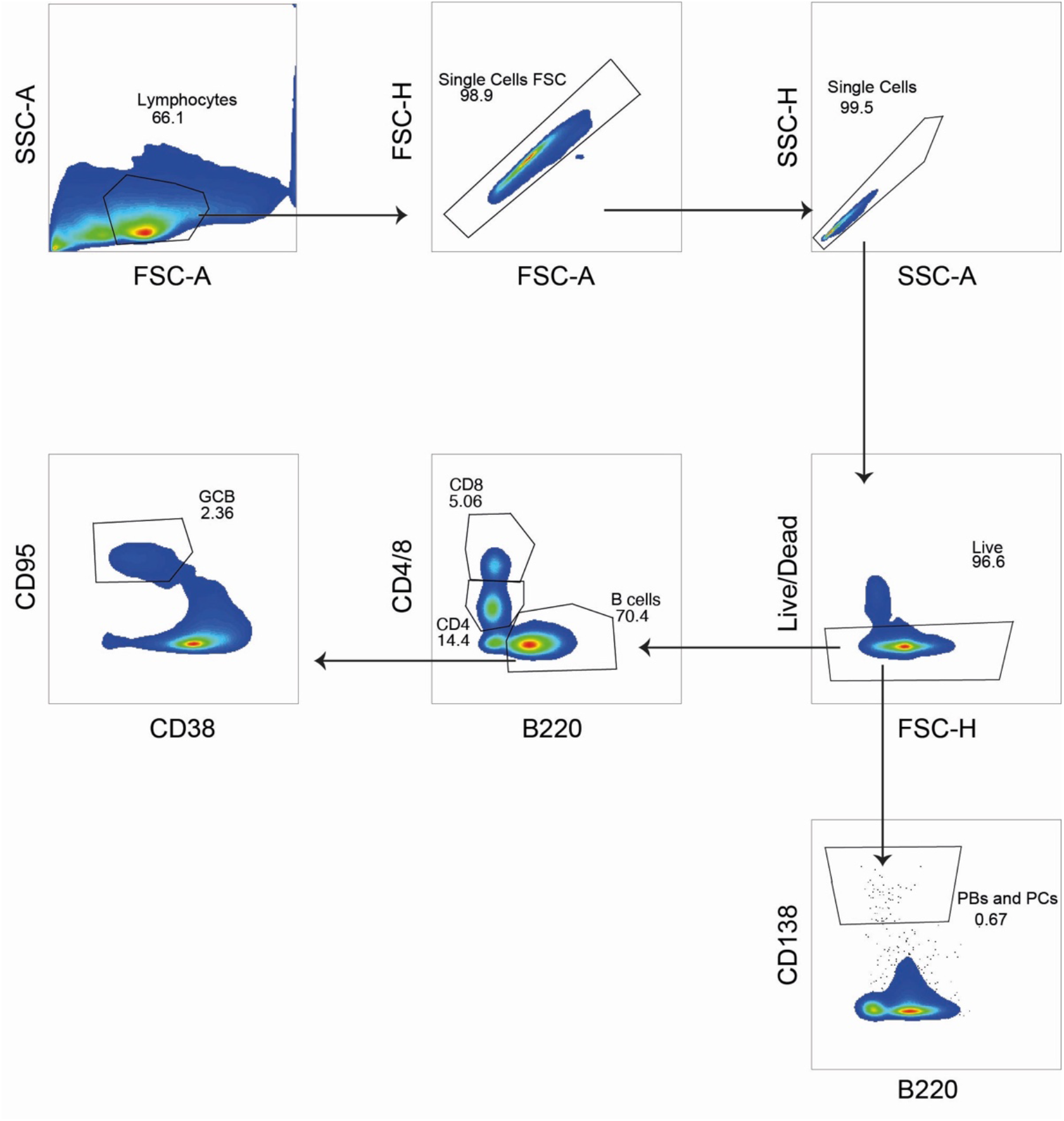
Gating strategy for the mixed bone-marrow chimera model cohort: Leukocytes were gated based on size (FSC-A) and granularity (SSC-A). Doublets were excluded with two singlet gates, first FSC-H vs FSC-A, and then SSC-H vs. SSC-A. Dead cells were excluded. B cells were selected from the live gate, from which GCBs were selected based on CD95 expression and the absence of CD38 expression. PBs and PCs were selected from the live gate based on CD138 expression.

### The extrafollicular pathway is sufficient to drive early hallmarks of autoimmunity in a second, independent lupus model

To further evaluate the robustness of our central observation that extrafollicular responses were sufficient to compensate or could even exacerbate autoimmune progression, we set out to test the effect of a complete GC block in a second, independent lupus model, 564Igi. The 564Igi model, which we also used as the driver compartment in the mixed chimeras, is a B cell receptor knock-in model, derived from a B cell clone isolated from an autoreactive F1 hybrid cross of Swiss Inbred (SWR) and New Zealand Black (NZB) mice (Berland et al., 2006; Gavalchin et al., 1985). The clone was originally screened for reactivity with ssDNA, but has been found to react more generally with ribonucleic acids and several ribonuclear proteins (Degn et al., 2017b), a polyreactivity likely owing to a high cationic pI in its complementarity determining region (Gavalchin et al., 1987). 564Igi mice present with spontaneous GCs and robust levels of circulating anti-DNA antibodies, but do not develop more severe hallmarks of disease until later in life (at least 9-12 months of age). We crossed the 564Igi line with Aicda-Cre Bcl6^flx/flx^ mice, and backcrossed the heavy and light chain, as well as Bcl6^flx^ to homozygosity, retaining hemiparental Aicda-Cre hemizygosity. Hence, Aicda-Cre positive vs. negative offspring, which were invariably 564Igi and Bcl6^flx^ homozygous, could be compared (**Fig. 6A and S6A**). We analyzed this offspring either at adolescence (13-16 weeks) (**Figure 6**) or in early adulthood (17-23 weeks of age) (**Figure S6**). At the earlier time point, the two groups had comparable levels of anti-dsDNA of IgG2a isotype (**Fig. 6B**), whereas GC block 564Igi mice presented with slightly elevated levels of anti-dsDNA IgG2b (**Fig. 6C**), although this difference did not reach statistical significance. Total anti-dsDNA levels were comparable between groups (**Fig. 6D**). Total IgG1 was elevated in GC-sufficient littermates (**Fig. 6E**), whereas total IgG2a (**Fig. 6F**) and IgG3 (**Fig. 6G**) were on par. By flow cytometry (**Fig. S7**), we verified that main lymphocyte subsets were comparable between groups across all tissues (**Fig. 6H-J**). Furthermore, the frequency of idiotype positive cells, i.e., cells carrying the knock-in B cell receptor (identified by staining with the anti-idiotype antibody 9D11) was comparable between groups (**Fig. 6K**). We verified that GCs were indeed efficiently blocked in GC block mice (**Fig. 6L**), but despite this, overall PB/PC frequencies (**Fig. 6M**) and PB frequencies (**Fig. 6N**) across tissues were not different between the two groups. PC frequencies were similar in IngLN and MesLN, and slightly elevated in the spleen of GC block mice (**Fig. 6O**). Thus, overall, the GC block did not prevent any of the autoimmune read-outs. This was also the case at the later time point, where no gross differences were observed (**Fig. S6**).

**Figure 6.**
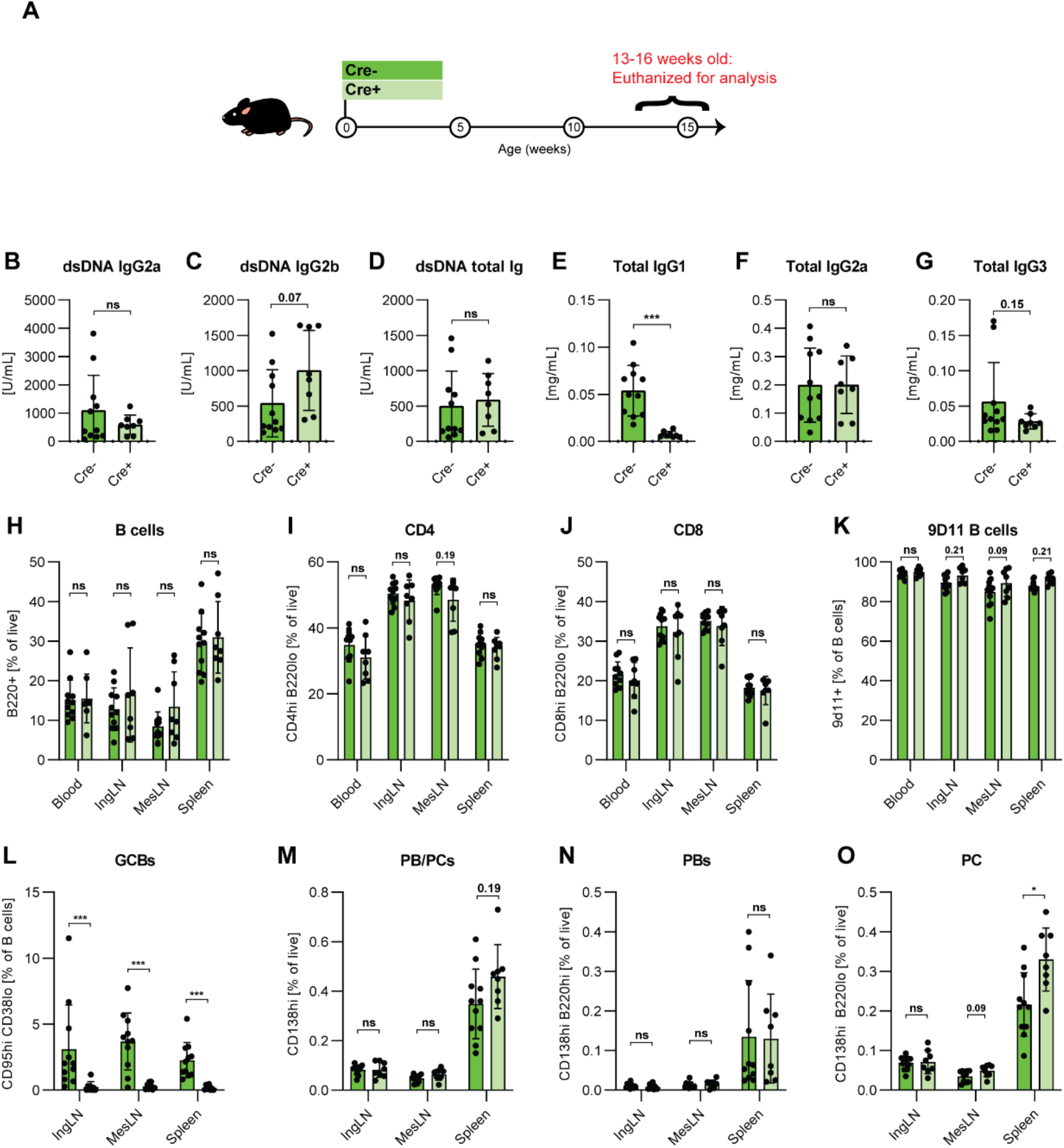
GC block causes increased PB/PCs levels in adolescent 564Igi mice. (**A**) Schematic overview of experimental setup: 564Igi-Bcl-6^flx/flx^ (Cre−, dark green) and 564Igi-Aicda-Cre Bcl-6^flx/flx^ (Cre+, light green) mice. (**B**) Anti-dsDNA IgG2a. (**C**) Anti-dsDNA IgG2b. (**D**) Anti-dsDNA total Ig. (**E**) Total IgG1. (**F**) Total IgG2a. (**G**) Total IgG3. Unpaired t-test or Mann-Whitney’s test was used was used to analyze TRIFMA data. (**H**) Flow cytometry analyses of B cell frequencies (B220^+^ CD4^-^CD8^-^of live, singlet lymphocytes) in blood, IngLN, MesLN, and spleen. (**I**) As H, but CD4 T cells (CD4^+^ B220^-^of live, singlet lymphocytes). (**J**) As H, but CD8 T cells (CD8^+^ B220^-^of live, singlet lymphocytes). (**K**) As H, but 9D11 B cell frequencies (9D11^+^ of B cells). (**L**) As H, but GCB frequencies (CD95^hi^ CD38^lo^ of B cells). (**M**) As H, but PB and PC frequencies (CD138^hi^ of live, singlet leukocytes), (**N**) As H, but PC frequencies (CD138^hi^ B220^lo^ of live, singlet lymphocytes), (**O**) As H, but PB frequencies (CD138^hi^ B220^hi^ of live, singlet lymphocytes). Data are pooled from two independent experiments. Bar graphs show mean ± SD. Two-way ANOVA with Holm-Sidak’s post hoc test was used to analyze the data. ns = p≥0.05, * = p<0.05, ** = p<0.01, *** = p<0.001.

Taken together, this demonstrated that the GC block failed to curb the main hallmarks of autoimmunity presented by the 564Igi model, further corroborating our findings from the Resiquimod model.

**Figure S6.**
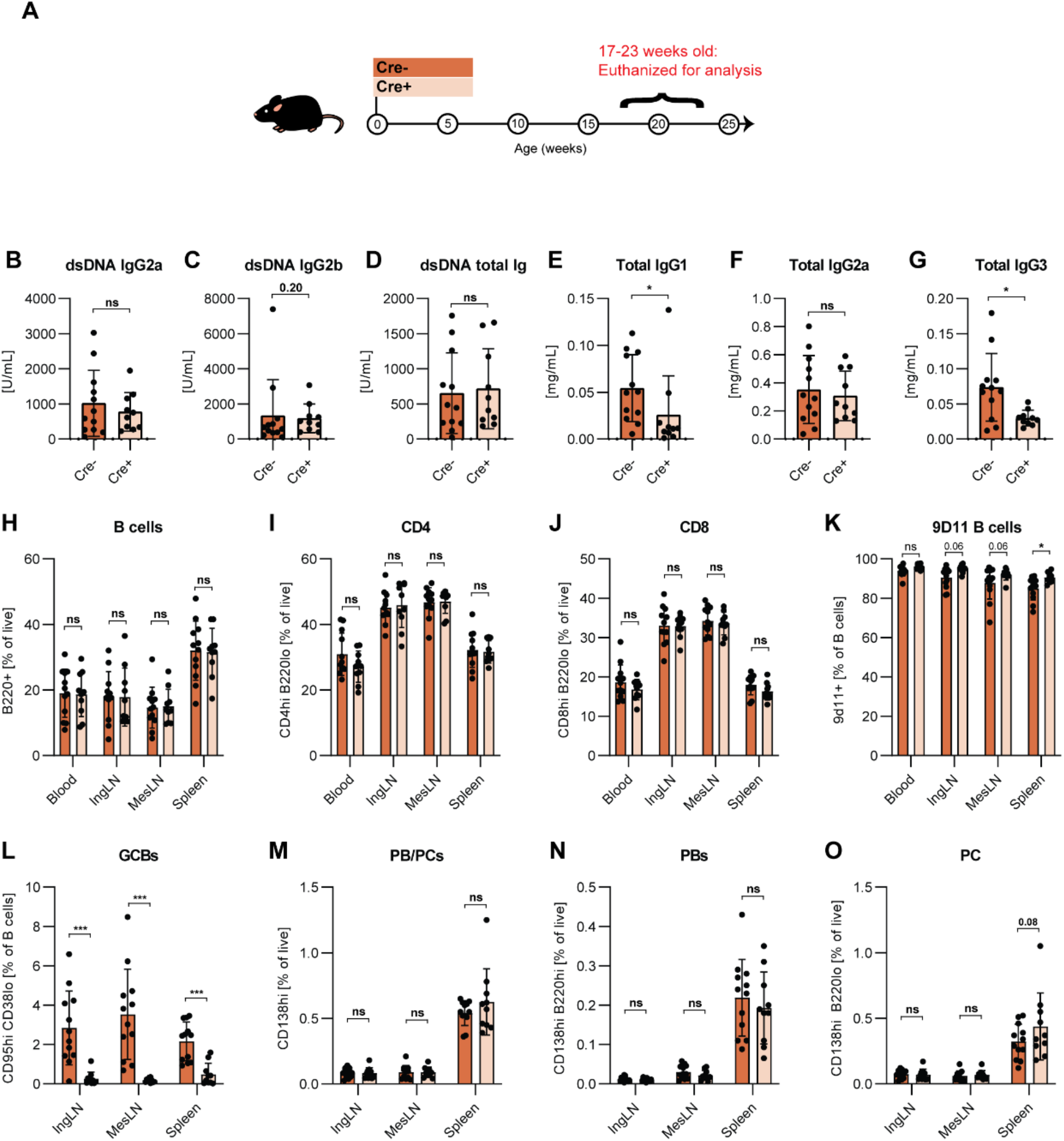
GC block does not ameliorate autoimmune phenotype in adult 564Igi mice. (**A**) Schematic overview of experimental setup: 564Igi-Bcl-6^flx/flx^ (Cre−, dark red) and 564Igi-Aicda-Cre Bcl-6^flx/flx^ (Cre+, light red). (**B**) Anti-dsDNA IgG2a. (**C**) Anti-dsDNA IgG2b. (**D**) Anti-dsDNA total Ig. (**E**) Total IgG1. (**F**) Total IgG2a. (**G**) Total IgG3. Unpaired t-test or Mann-Whitney’s test was used was used to analyze TRIFMA data. (**H**) Flow cytometry analyses of B cell frequencies (B220^+^ CD4^-^CD8^-^of live, singlet lymphocytes) in blood, IngLN, MesLN, and spleen. (**I**) As H, but CD4 T cells (CD4^+^ B220^-^of live, singlet lymphocytes). (**J**) As H, but CD8 T cells (CD8^+^ B220^-^of live, singlet lymphocytes). (**K**) As H, but 9D11 B cell frequencies (9D11^+^ of B cells). (**L**) As H, but GCB frequencies (CD95^hi^ CD38^lo^ of B cells). (**M**) As H, but PB and PC frequencies (CD138^hi^ of live, singlet leukocytes), (**N**) As H, but PC frequencies (CD138^hi^ B220^lo^ of live, singlet lymphocytes), (**O**) As H, but PB frequencies (CD138^hi^ B220^hi^ of live, singlet lymphocytes). Data are pooled from two independent experiments. Bar graphs show mean ± SD. Two-way ANOVA with Holm-Sidak’s post hoc test was used to analyze the data. ns = p≥0.05, * = p<0.05, ** = p<0.01, *** = p<0.001.

**Supplementary Figure 7.**
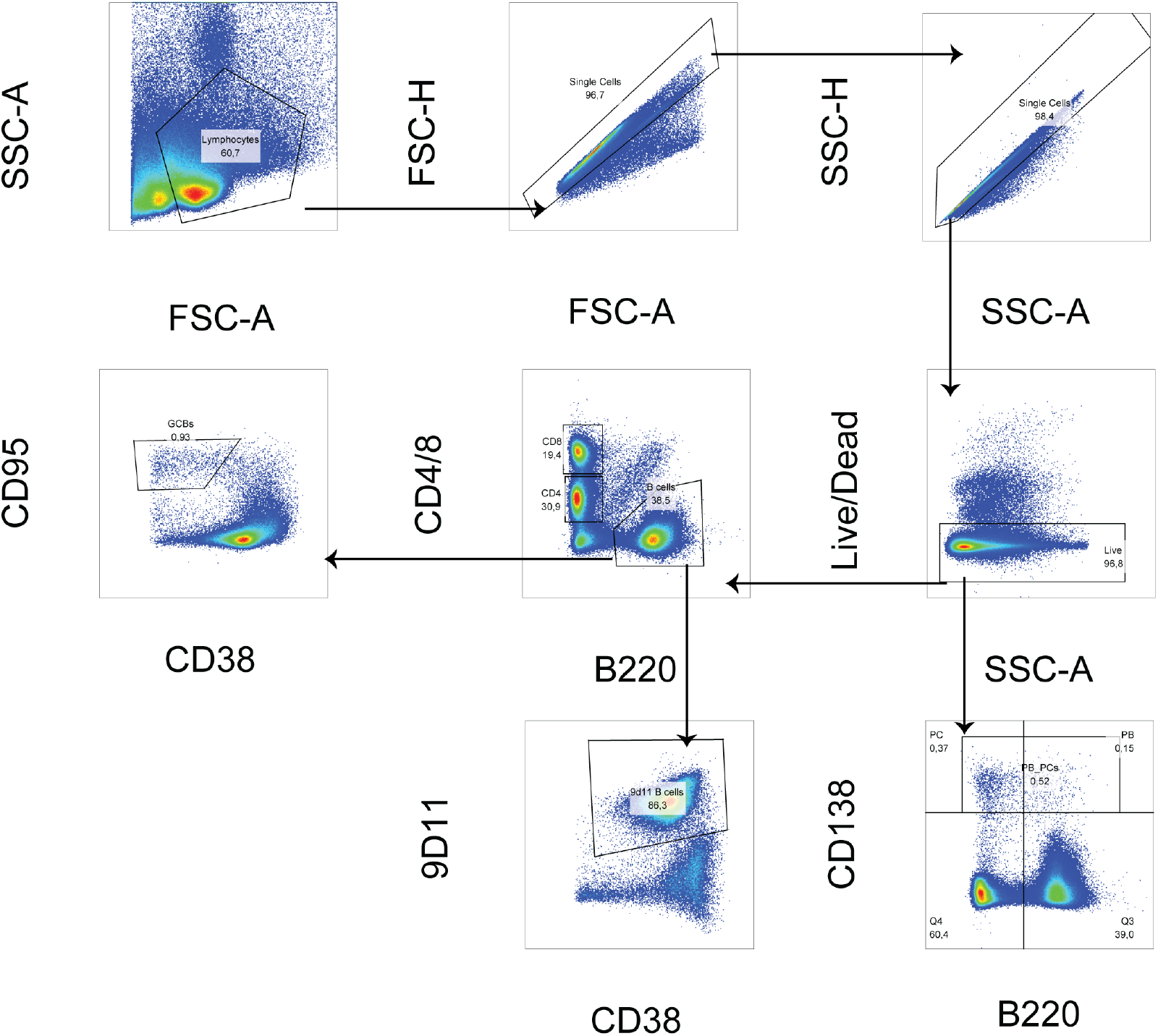
Gating strategy for the 564Igi model cohort: Lymphocytes were gated based on size (FSC-A) and granularity (SSC-A). Doublets were excluded with two singlet gates, first FSC-H vs FSC-A, and then SSC-H vs. SSC-A. Dead cells were excluded. B cells were selected from the live gate, from which GCBs were selected based on CD95 expression and the absence of CD38 expression. Idiotype (9D11) positive cells were selected from the B cell gate based on 9D11^hi^ expression. PBs and PCs were selected from the live gate based on CD138 and B220 expression.

## Discussion

GCs are believed to be the nexus of autoreactive responses in a range of autoimmune diseases. Due to their role in potent antibody responses, memory generation, and long-lived PC formation, there has been extensive interest in targeting GCs in autoimmune disease. The strategy has proven useful in autoimmune models, but due to off-target effects, did not initially progress through clinical trials (Degn et al., 2017a; Karnell et al., 2019). Considerable efforts have, however, been aimed at circumventing the off-target effects to bring this strategy to market (Espie et al., 2020; Karnell et al., 2019). This notwithstanding, a recent study reported that extrafollicular B cell differentiation into short-lived antibody-forming cells is a key mechanism of anti-DNA autoreactivity (Soni et al., 2020), and it has been argued that more attention should be paid to the non-GC responses, as these may play a critical role in humoral immunity in both mice and men (Jenks et al., 2019).

Here, we took an unbiased approach and asked to what extent a genetic block of GCs would ameliorate autoreactive manifestations in a lupus-like disease model. To our surprise, we found blocking GCs did not lessen autoreactive manifestations, but in some cases worsened these. Upon autoimmune induction, we observed a trend towards an increase in anti-dsDNA antibodies of the IgG2c isotype (**Fig. 1C**) and total IgG2c antibody (**Fig. 1D**) in GC block mice, compared to WT. These changes were also mirrored by a significantly higher frequency of PB/PCs in the spleens of GC block mice, despite a total absence of GCs. In agreement with this, we observed robust deposition of immune complexes in the kidney glomeruli of GC block mice, at least on par with that of GC sufficient controls (**Fig. 3**). This indicated that the extrafollicular pathway could compensate, and in some cases even augment, the autoreactive response. We corroborated this central finding in a second, independent model (**Fig. 6**), confirming that the observed sufficiency of the extrafollicular response was not model-specific.

To understand the B cell intrinsic effect of a GC block, we leveraged an induced GC B cell culture system. It has previously been noted that Bcl-6 expression can inhibit apoptosis in numerous cell types including (GC) B cells (Kumagai et al., 1999; Kurosu et al., 2003; Phan and Dalla-Favera, 2004). Yet, contrary to expectations, GC block B cells did not display a significantly increased propensity to undergo apoptosis (**Fig. 4H and L**), rather, they much more readily underwent terminal differentiation to PCs, and had a dramatically reduced capacity to expand compared to their wild type counterparts (**Fig. 4**). However, this agreed well with the established cross-regulation between Bcl-6 and Blimp-1, the master regulator of the PC fate, also known as Prdm1 (Vasanwala et al., 2002). Although the increased propensity for terminal PC differentiation was, in principle, well in line with our *in vivo* observations, the lack of proliferative capacity was at the same time at odds with the dramatic PC output in the mice harboring a GC block in B cells. This suggested that the PC differentiation process in mice displaying a global GC block in B cells might be dysregulated, potentially as a consequence of absence of GC-derived antibody feedback, as previously suggested for GC B cells (Zhang et al., 2013). To address this possibility, we asked whether B cells with a GC block would be precluded from contributing to the PC pool in a GC sufficient environment. Our findings demonstrated that this was not the case, although the relative contribution of GC block B cells to the PC pool was smaller than that of GC sufficient B cells (**Fig. 5**). However, given their inability to expand in GCs, the magnitude of the contribution of GC block B cells to the PC compartment in direct competition with GC-sufficient B cells was remarkable. In the infectious setting, an early wave of extrafollicular PCs is crucial for the initial antibody response. However, most PCs produced by the extrafollicular response undergo apoptosis within a matter of days, and the global response becomes dominated by GC-derived responses. In the chronic autoreactive setting, however, the continuous fueling of the autoimmune process may continually renew this population.

We may speculate that the somewhat lower contribution of the extrafollicular PC compartment in the mixed chimera model compared to that of the R848 model could be a consequence of the markedly different time scales of the two experiments: the mixed chimeras were analyzed 13 weeks post reconstitution, whereas the R848 mice were analyzed 4.5 weeks after commencement of treatment. This could be important, because at this point it remains unclear whether the GC responses observed in our models contribute a qualitatively different response to the autoimmune progression, e.g., through production of memory B cells and long-lived PCs that may perpetuate and dominate the chronic response over longer periods of time. By extension, the GC pathway may differentially allow epitope spreading and inclusion of alternative antigens over time, as seen in human SLE patients (Arbuckle et al., 2003). At least, it seems plausible that the longer the autoimmune process has persisted, the more the long-lived GC responses and their derived memory output come to dominate the process, and conversely, the short-lived extrafollicular responses may govern the early stages of the response and, as previously suggested, the very early break-of-tolerance driven by nucleic acid-containing antigens (Soni et al., 2020; Sweet et al., 2011). Such considerations notwithstanding, our findings in the second model, 564Igi, analyzed at 13-16 and 17-23 weeks of age, suggest that even after a chronic response of 5-6 months in duration, the extrafollicular responses are not outpaced by GCs (**Fig. 6 and S6**).

Interestingly, due to its more potent nature, The GC reaction is believed to be subject to a much higher level of control, through a continued requirement for linked recognition in successive rounds of diversity generation. Furthermore, a specialized subset of Tregs, T_FR_s, exert a dominant negative level of control on the GC reaction (Fahlquist Hagert and Degn, 2020). Hence it may be that the extrafollicular pathway in essence represents an evolutionary ‘backdoor to autoimmunity’, unguarded due to its relative insignificance in terms of high-quality, affinity-matured, and memory-inducing antibody responses. In this context, it is fortunate that current CD40L targeting strategies block both the GC and extrafollicular response. However, we suggest that future efforts should be aimed at further elucidating the relative contributions of the extrafollicular and GC pathways. We may speculate that specific targeting of the extrafollicular pathway would be a superior strategy, as it would preferentially block the low quality and poorly controlled responses driving autoimmune progression, while leaving intact the more stringently controlled and high-quality responses that provide protection against infectious agents. Unfortunately, there is much more limited knowledge regarding the biology of the extrafollicular responses, and no transgenic or pharmacologic strategy allowing specific blockade of this pathway exists, making it difficult to evaluate in animal models.

In summary, our findings here demonstrate that a complete or partial block of the GC pathway is insufficient to curb autoreactive PC differentiation and might in fact exacerbate the autoimmune progression. The GC commitment is controlled by the expression level of the master transcriptional repressor, Bcl-6 (Robinson et al., 2020), which regulates the GC fates across GC B cells, T_FH_ cells and T_FR_ cells. Interestingly, in the context of the COVID-19 pandemic, it has been observed that Bcl-6^+^ GC B cells and Bcl-6^+^ T_FH_ cells are markedly diminished in SARS-CoV-2 infection (Kaneko et al., 2020). It has also been found that critically ill SARS-CoV-2 patients display hallmarks of extrafollicular B cell activation and shared B cell repertoire features previously described in autoimmune settings (Knight et al., 2021; Woodruff et al., 2020). This further highlights the potential link between aberrant extrafollicular responses and autoimmune manifestations.

## Acknowledgements

We thank Hanne Sidelmann for her contributions to immunofluorescence and immunohistochemical staining of kidneys. We would like to acknowledge the AU FACS Core facility for their support and feedback on data related to flow cytometry. We also thank the BioImaging Core at Health for support with microscopy.

## References

Akkaya, M., B. Akkaya, A.S. Kim, P. Miozzo, H. Sohn, M. Pena, A.S. Roesler, B.P. Theall, T. Henke, J. Kabat, J. Lu, D.W. Dorward, E. Dahlstrom, J. Skinner, L.H. Miller, and S.K. Pierce. 2018. Toll-like receptor 9 antagonizes antibody affinity maturation. Nat Immunol 19:255–266.

Arbuckle, M.R., M.T. McClain, M.V. Rubertone, R.H. Scofield, G.J. Dennis, J.A. James, and J.B. Harley. 2003. Development of autoantibodies before the clinical onset of systemic lupus erythematosus. N Engl J Med 349:1526–1533.

Berland, R., L. Fernandez, E. Kari, J.H. Han, I. Lomakin, S. Akira, H.H. Wortis, J.F. Kearney, A.A. Ucci, and T. Imanishi-Kari. 2006. Toll-like receptor 7-dependent loss of B cell tolerance in pathogenic autoantibody knockin mice. Immunity 25:429–440.

Cornaby, C., L. Gibbons, V. Mayhew, C.S. Sloan, A. Welling, and B.D. Poole. 2015. B cell epitope spreading: mechanisms and contribution to autoimmune diseases. Immunol Lett 163:56–68.

Degn, S.E., C.E. van der Poel, and M.C. Carroll. 2017a. Targeting autoreactive germinal centers to curb autoimmunity. Oncotarget 8:90624–90625.

Degn, S.E., C.E. van der Poel, D.J. Firl, B. Ayoglu, F.A. Al Qureshah, G. Bajic, L. Mesin, C.A. Reynaud, J.C. Weill, P.J. Utz, G.D. Victora, and M.C. Carroll. 2017b. Clonal Evolution of Autoreactive Germinal Centers. Cell 170:913–926 e919.

Domeier, P.P., S.L. Schell, and Z.S. Rahman. 2017. Spontaneous germinal centers and autoimmunity. Autoimmunity 50:4–18.

Elsner, R.A., and M.J. Shlomchik. 2020. Germinal Center and Extrafollicular B Cell Responses in Vaccination, Immunity, and Autoimmunity. Immunity 53:1136–1150.

Espie, P., Y. He, P. Koo, D. Sickert, C. Dupuy, E. Chokote, R. Schuler, H. Mergentaler, J. Ristov, J. Milojevic, A. Verles, A. Groenewegen, A. Auger, A. Avrameas, M. Rotte, L. Colin, C.S. Tomek, M. Hernandez-Illas, J.S. Rush, and P. Gergely. 2020. First-in-human clinical trial to assess pharmacokinetics, pharmacodynamics, safety, and tolerability of iscalimab, an anti-CD40 monoclonal antibody. Am J Transplant 20:463–473.

Fahlquist Hagert, C., and S.E. Degn. 2020. T follicular regulatory cells: Guardians of the germinal centre? Scand J Immunol 92:e12942.

Gasparotto, M., M. Gatto, V. Binda, A. Doria, and G. Moroni. 2020. Lupus nephritis: clinical presentations and outcomes in the 21st century. Rheumatology (Oxford) 59:v39–v51.

Gavalchin, J., J.A. Nicklas, J.W. Eastcott, M.P. Madaio, B.D. Stollar, R.S. Schwartz, and S.K. Datta. 1985. Lupus prone (SWR x NZB)F1 mice produce potentially nephritogenic autoantibodies inherited from the normal SWR parent. J Immunol 134:885–894.

Gavalchin, J., R.A. Seder, and S.K. Datta. 1987. The NZB X SWR model of lupus nephritis. I. Cross-reactive idiotypes of monoclonal anti-DNA antibodies in relation to antigenic specificity, charge, and allotype. Identification of interconnected idiotype families inherited from the normal SWR and the autoimmune NZB parents. J Immunol 138:128–137.

Green, K., T.R. Wittenborn, C. Fahlquist-Hagert, E. Terczynska-Dyla, N. van Campen, L. Jensen, L. Reinert, R. Hartmann, S.R. Paludan, and S.E. Degn. 2021. B Cell Intrinsic STING Signaling Is Not Required for Autoreactive Germinal Center Participation. Front Immunol 12:782558.

Hollister, K., S. Kusam, H. Wu, N. Clegg, A. Mondal, D.V. Sawant, and A.L. Dent. 2013. Insights into the role of Bcl6 in follicular Th cells using a new conditional mutant mouse model. J Immunol 191:3705–3711.

Jenks, S.A., K.S. Cashman, M.C. Woodruff, F.E. Lee, and I. Sanz. 2019. Extrafollicular responses in humans and SLE. Immunol Rev 288:136–148.

Juul-Madsen, K., A. Troldborg, T.R. Wittenborn, M.G. Axelsen, H. Zhao, L.H. Klausen, S. Luecke, S.R. Paludan, K. Stengaard-Pedersen, M. Dong, H.J. Moller, S. Thiel, H. Jensen, P. Schuck, D.S. Sutherland, S.E. Degn, and T. Vorup-Jensen. 2021. Characterization of DNA-protein complexes by nanoparticle tracking analysis and their association with systemic lupus erythematosus. Proc Natl Acad Sci U S A 118:

Kaneko, N., H.H. Kuo, J. Boucau, J.R. Farmer, H. Allard-Chamard, V.S. Mahajan, A. Piechocka-Trocha, K. Lefteri, M. Osborn, J. Bals, Y.C. Bartsch, N. Bonheur, T.M. Caradonna, J. Chevalier, F. Chowdhury, T.J. Diefenbach, K. Einkauf, J. Fallon, J. Feldman, K.K. Finn, P. Garcia-Broncano, C.A. Hartana, B.M. Hauser, C. Jiang, P. Kaplonek, M. Karpell, E.C. Koscher, X. Lian, H. Liu, J. Liu, N.L. Ly, A.R. Michell, Y. Rassadkina, K. Seiger, L. Sessa, S. Shin, N. Singh, W. Sun, X. Sun, H.J. Ticheli, M.T. Waring, A.L. Zhu, G. Alter, J.Z. Li, D. Lingwood, A.G. Schmidt, M. Lichterfeld, B.D. Walker, X.G. Yu, R.F. Padera, Jr., S. Pillai, and G. Massachusetts Consortium on Pathogen Readiness Specimen Working. 2020. Loss of Bcl-6-Expressing T Follicular Helper Cells and Germinal Centers in COVID-19. Cell 183:143–157 e113.

Karnell, J.L., M. Albulescu, S. Drabic, L. Wang, R. Moate, M. Baca, V. Oganesyan, M. Gunsior, T. Thisted, L. Yan, J. Li, X. Xiong, S.C. Eck, M. de Los Reyes, I. Yusuf, K. Streicher, U. Muller-Ladner, D. Howe, R. Ettinger, R. Herbst, and J. Drappa. 2019. A CD40L-targeting protein reduces autoantibodies and improves disease activity in patients with autoimmunity. Sci Transl Med 11:

Knight, J.S., R. Caricchio, J.L. Casanova, A.J. Combes, B. Diamond, S.E. Fox, D.A. Hanauer, J.A. James, Y. Kanthi, V. Ladd, P. Mehta, A.M. Ring, I. Sanz, C. Selmi, R.P. Tracy, P.J. Utz, C.A. Wagner, J.Y. Wang, and W.J. McCune. 2021. The intersection of COVID-19 and autoimmunity. J Clin Invest 131:

Kumagai, T., T. Miki, M. Kikuchi, T. Fukuda, N. Miyasaka, R. Kamiyama, and S. Hirosawa. 1999. The proto-oncogene Bc16 inhibits apoptotic cell death in differentiation-induced mouse myogenic cells. Oncogene 18:467–475.

Kuraoka, M., A.G. Schmidt, T. Nojima, F. Feng, A. Watanabe, D. Kitamura, S.C. Harrison, T.B. Kepler, and G. Kelsoe. 2016. Complex Antigens Drive Permissive Clonal Selection in Germinal Centers. Immunity 44:542–552.

Kurosu, T., T. Fukuda, T. Miki, and O. Miura. 2003. BCL6 overexpression prevents increase in reactive oxygen species and inhibits apoptosis induced by chemotherapeutic reagents in B-cell lymphoma cells. Oncogene 22:4459–4468.

Kwon, K., C. Hutter, Q. Sun, I. Bilic, C. Cobaleda, S. Malin, and M. Busslinger. 2008. Instructive role of the transcription factor E2A in early B lymphopoiesis and germinal center B cell development. Immunity 28:751–762.

Lau, C.M., C. Broughton, A.S. Tabor, S. Akira, R.A. Flavell, M.J. Mamula, S.R. Christensen, M.J. Shlomchik, G.A. Viglianti, I.R. Rifkin, and A. Marshak-Rothstein. 2005. RNA-associated autoantigens activate B cells by combined B cell antigen receptor/Toll-like receptor 7 engagement. J Exp Med 202:1171–1177.

Leadbetter, E.A., I.R. Rifkin, A.M. Hohlbaum, B.C. Beaudette, M.J. Shlomchik, and A. Marshak-Rothstein. 2002. Chromatin-IgG complexes activate B cells by dual engagement of IgM and Toll-like receptors. Nature 416:603–607.

Lee, S.K., R.J. Rigby, D. Zotos, L.M. Tsai, S. Kawamoto, J.L. Marshall, R.R. Ramiscal, T.D. Chan, D. Gatto, R. Brink, D. Yu, S. Fagarasan, D.M. Tarlinton, A.F. Cunningham, and C.G. Vinuesa. 2011. B cell priming for extrafollicular antibody responses requires Bcl-6 expression by T cells. J Exp Med 208:1377–1388.

Luzina, I.G., S.P. Atamas, C.E. Storrer, L.C. daSilva, G. Kelsoe, J.C. Papadimitriou, and B.S. Handwerger. 2001. Spontaneous formation of germinal centers in autoimmune mice. J Leukoc Biol 70:578–584.

Nojima, T., K. Haniuda, T. Moutai, M. Matsudaira, S. Mizokawa, I. Shiratori, T. Azuma, and D. Kitamura. 2011. In-vitro derived germinal centre B cells differentially generate memory B or plasma cells in vivo. Nat Commun 2:465.

Paus, D., T.G. Phan, T.D. Chan, S. Gardam, A. Basten, and R. Brink. 2006. Antigen recognition strength regulates the choice between extrafollicular plasma cell and germinal center B cell differentiation. J Exp Med 203:1081–1091.

Phan, R.T., and R. Dalla-Favera. 2004. The BCL6 proto-oncogene suppresses p53 expression in germinal-centre B cells. Nature 432:635–639.

Psianou, K., I. Panagoulias, A.D. Papanastasiou, A.L. de Lastic, M. Rodi, P.I. Spantidea, S.E. Degn, P. Georgiou, and A. Mouzaki. 2018. Clinical and immunological parameters of Sjogren’s syndrome. Autoimmun Rev 17:1053–1064.

Rahman, A., and D.A. Isenberg. 2008. Systemic lupus erythematosus. N Engl J Med 358:929–939.

Robinson, M.J., Z. Ding, C. Pitt, E.J. Brodie, I. Quast, D.M. Tarlinton, and D. Zotos. 2020. The Amount of BCL6 in B Cells Shortly after Antigen Engagement Determines Their Representation in Subsequent Germinal Centers. Cell Rep 30:1530–1541 e1534.

Roco, J.A., L. Mesin, S.C. Binder, C. Nefzger, P. Gonzalez-Figueroa, P.F. Canete, J. Ellyard, Q. Shen, P.A. Robert, J. Cappello, H. Vohra, Y. Zhang, C.R. Nowosad, A. Schiepers, L.M. Corcoran, K.M. Toellner, J.M. Polo, M. Meyer-Hermann, G.D. Victora, and C.G. Vinuesa. 2019. Class-Switch Recombination Occurs Infrequently in Germinal Centers. Immunity 51:337–350 e337.

Soni, C., O.A. Perez, W.N. Voss, J.N. Pucella, L. Serpas, J. Mehl, K.L. Ching, J. Goike, G. Georgiou, G.C. Ippolito, V. Sisirak, and B. Reizis. 2020. Plasmacytoid Dendritic Cells and Type I Interferon Promote Extrafollicular B Cell Responses to Extracellular Self-DNA. Immunity 52:1022–1038 e1027.

Sweet, R.A., S.R. Christensen, M.L. Harris, J. Shupe, J.L. Sutherland, and M.J. Shlomchik. 2010. A new site-directed transgenic rheumatoid factor mouse model demonstrates extrafollicular class switch and plasmablast formation. Autoimmunity 43:607–618.

Sweet, R.A., M.L. Ols, J.L. Cullen, A.V. Milam, H. Yagita, and M.J. Shlomchik. 2011. Facultative role for T cells in extrafollicular Toll-like receptor-dependent autoreactive B-cell responses in vivo. Proc Natl Acad Sci U S A 108:7932–7937.

Toellner, K.M., A. Gulbranson-Judge, D.R. Taylor, D.M. Sze, and I.C. MacLennan. 1996. Immunoglobulin switch transcript production in vivo related to the site and time of antigen-specific B cell activation. J Exp Med 183:2303–2312.

Vasanwala, F.H., S. Kusam, L.M. Toney, and A.L. Dent. 2002. Repression of AP-1 function: a mechanism for the regulation of Blimp-1 expression and B lymphocyte differentiation by the B cell lymphoma-6 protooncogene. J Immunol 169:1922–1929.

Victora, G.D., and M.C. Nussenzweig. 2022. Germinal Centers. Annu Rev Immunol

Weening, J.J., V.D. D’Agati, M.M. Schwartz, S.V. Seshan, C.E. Alpers, G.B. Appel, J.E. Balow, J.A. Bruijn, T. Cook, F. Ferrario, A.B. Fogo, E.M. Ginzler, L. Hebert, G. Hill, P. Hill, J.C. Jennette, N.C. Kong, P. Lesavre, M. Lockshin, L.M. Looi, H. Makino, L.A. Moura, and M. Nagata. 2004. The classification of glomerulonephritis in systemic lupus erythematosus revisited. J Am Soc Nephrol 15:241–250.

William, J., C. Euler, S. Christensen, and M.J. Shlomchik. 2002. Evolution of autoantibody responses via somatic hypermutation outside of germinal centers. Science 297:2066–2070.

Wittenborn, T.R., C. Fahlquist Hagert, A. Ferapontov, S. Fonager, L. Jensen, G. Winther, and S.E. Degn. 2021. Comparison of gamma and x-ray irradiation for myeloablation and establishment of normal and autoimmune syngeneic bone marrow chimeras. PLoS One 16:e0247501.

Woodruff, M.C., R.P. Ramonell, D.C. Nguyen, K.S. Cashman, A.S. Saini, N.S. Haddad, A.M. Ley, S. Kyu, J.C. Howell, T. Ozturk, S. Lee, N. Suryadevara, J.B. Case, R. Bugrovsky, W. Chen, J. Estrada, A. Morrison-Porter, A. Derrico, F.A. Anam, M. Sharma, H.M. Wu, S.N. Le, S.A. Jenks, C.M. Tipton, B. Staitieh, J.L. Daiss, E. Ghosn, M.S. Diamond, R.H. Carnahan, J.E. Crowe, Jr., W.T. Hu, F.E. Lee, and I. Sanz. 2020. Extrafollicular B cell responses correlate with neutralizing antibodies and morbidity in COVID-19. Nat Immunol 21:1506–1516.

Yokogawa, M., M. Takaishi, K. Nakajima, R. Kamijima, C. Fujimoto, S. Kataoka, Y. Terada, and S. Sano. 2014. Epicutaneous application of toll-like receptor 7 agonists leads to systemic autoimmunity in wild-type mice: a new model of systemic Lupus erythematosus. Arthritis Rheumatol 66:694–706.

Zhang, Y., M. Meyer-Hermann, L.A. George, M.T. Figge, M. Khan, M. Goodall, S.P. Young, A. Reynolds, F. Falciani, A. Waisman, C.A. Notley, M.R. Ehrenstein, M. Kosco-Vilbois, and K.M. Toellner. 2013. Germinal center B cells govern their own fate via antibody feedback. J Exp Med 210:457–464.

